# Tensor Decomposition of Stimulated Monocyte and Macrophage Gene Expression Profiles Identifies Neurodegenerative Disease-specific *Trans*-eQTLs

**DOI:** 10.1101/499509

**Authors:** Satesh Ramdhani, Elisa Navarro, Evan Udine, Brian M. Schilder, Madison Parks, Towfique Raj

**Affiliations:** Ronald M. Loeb Center for Alzheimer’s Disease, Icahn School of Medicine at Mount Sinai, New York, New York, USA; Department of Neuroscience and Friedman Brain Institute, Icahn School of Medicine at Mount Sinai, New York, New York, USA; Department of Genetics and Genomic Sciences, Icahn School of Medicine at Mount Sinai, New York, New York, USA

## Abstract

Recent human genetic studies suggest that cells of the innate immune system have a primary role in the pathogenesis of neurodegenerative diseases. However, the results from these studies often do not elucidate how the genetic variants affect the biology of these cells to modulate disease risk. Here, we applied a tensor decomposition method to uncover disease-associated gene networks linked to distal genetic variation in stimulated human monocytes and macrophages gene expression profiles. We report robust evidence that some disease-associated genetic variants affect the expression of multiple genes in *trans*. These include a Parkinson’s disease locus influencing the expression of genes mediated by a protease that controls lysosomal function, and Alzheimer’s disease loci influencing the expression of genes involved in type 1 interferon signaling, myeloid phagocytosis, and complement cascade pathways. Overall, we uncover gene networks in induced innate immune cells linked to disease-associated genetic variants, which may help elucidate the underlying biology of disease.

## Introduction

Genome-wide associated studies (GWAS) have identified tens of thousands of common genetic variants associated with complex traits and diseases^1,2^. The vast majority of these genetic variants reside in non-coding regions^3^ potentially affecting the regulation of gene expression of local (*cis*) or distal (*trans*) genes. An extensive number of studies have characterized expression Quantitative Trait Loci (eQTL) across multiple tissues^4–6^ and cell types^7–10^ and in response to environmental stimuli^11,12^. Nevertheless, most eQTL studies in humans have been limited almost exclusively to *cis*-effects. The few studies that have investigated *trans*-eQTLs have identified few significant associations, and there has been little replication of significant associations across data sets. One limitation is that most of these studies used complex mixtures of different cell types (e.g., brain tissue, whole blood or peripheral blood mononuclear cells [PBMCs]) as RNA sources^13^, which may result in the failure to properly capture the activity of genetic variants in disease-relevant cell types. Another limitation is that these studies report *trans*-eQTLs by performing millions of Single Nucleotide Polymorphism (SNP)-by-gene tests, which can result in few significant associations due to the very stringent significance threshold imposed by multiple testing burden correction. Such SNP-by-gene *trans*-eQTL mapping also ignores the complex structure of gene networks. Recently, Hore, et al. ^14^ developed a method to decompose a tensor (or multi-dimensional array) of multi-tissue gene expression data to uncover gene networks and map these networks with genetic variants to detect *trans*-eQTLs. This and similar methods^15–17^ have been successful in mapping networks of genes regulated by genetic variants that would not have been uncovered via marginal SNP-by-gene *trans*-eQTL analysis.

Discovering such *trans*-eQTL networks may improve our understanding of the biological mechanisms underlying polygenic diseases such as Alzheimer’s disease and Parkinson’s disease. Both are neurodegenerative diseases which share pathological hallmarks such as neuronal loss, proteinopathy, mitochondrial dysfunction and reactive microglia (the innate immune cells of the brain)^18,19^, although the specific vulnerable cells differ between disorders. GWAS have identified over 30 and 41 loci, respectively, associated with Alzheimer’s disease ^20,21^ and Parkinson’s disease^22^. Nevertheless, these studies open new questions as they identify SNPs rather than genes, most of them localized in non-coding regions that are thought to modulate disease risk by influencing the expression of nearby or distal genes in a cell-type specific manner. Therefore, translating the effect of risk alleles to the genes and cell types involved in disease pathogenesis remain a challenge. Along this line, it should be noted that analyses of the human genetic data point to enrichment of Alzheimer’s disease and Parkinson’s disease risk alleles in genes, epigenetic annotations and cis-eQTL effects associated with myeloid cells of the innate immune system^7,23^. Animal models have also pointed to an altered innate immune system as a potential driver of these diseases^24–26^. While most research has focused on the contribution of microglia in neurodegeneration, there are reports suggesting a causal role of cellular and humoral components of the peripheral innate immune system in the pathogenesis and progression of these diseases. For example, parabiosis studies suggest that peripheral blood components may influence Alzheimer’s disease progression^27^. Whether peripheral blood monocytes may themselves be drivers of disease or are simply useful proxies for the infiltrating macrophages and/or resident microglia found at the sites of neuropathology is still unknown. We hypothesize that AD and PD risk alleles may modulate disease susceptibility by regulating the expression of a distal gene or set of genes in monocytes and macrophages (major cellular components of the innate immune system).

Here we used gene expression profiles from monocytes to identify novel *trans*-eQTLs and replicate those previously known. We show that about one-third of *trans* regulation is significantly mediated by the expression of *cis* genes. We identified *trans*-eQTLs that colocalize with disease-associated susceptibility loci. These include GWAS loci for coronary artery disease, body mass index, and cholesterol, all traits for which peripheral blood monocytes and macrophages are functionally relevant. More interestingly, our analysis also discovered biologically meaningful *trans*-eQTL networks for neurodegenerative diseases including genes in interferon and complement signaling, as well as lysosomal function linked to Alzheimer’s and Parkinson’s disease susceptibility loci, respectively. These results highlight context-specific *trans*-eQTL networks in support of a role for myeloid cells of the innate immune system as key modifiers of neurodegenerative disease risk.

## Results

### Identification of gene networks in stimulated monocytes and macrophages

We used Sparse Decomposition of Arrays (SDA)^14^ to construct gene networks by decomposing a multi-array set of gene expression measurements into latent components (see Methods). Each of these components consists of scores that indicate the relative contribution of each individual, gene, and cell or stimulation activity scores to gene networks (Fig. 1). The individual scores are the magnitude of the effect for each component across individuals. We use the individual scores as phenotypes in genome-wide *trans*-eQTL analysis to identify common genetic variants that drive each component. The gene scores (or gene loadings) allow to infer more clearly which genes are involved in each component. SDA is developed in a Bayesian framework and use ‘spike and slab’ prior^28^ to allow gene scores of each component to have unique level of sparsity. Finally, the cell or stimulation specificity scores indicate the activity of the component for each inflammatory stimuli or cell types.

**Figure 1:**
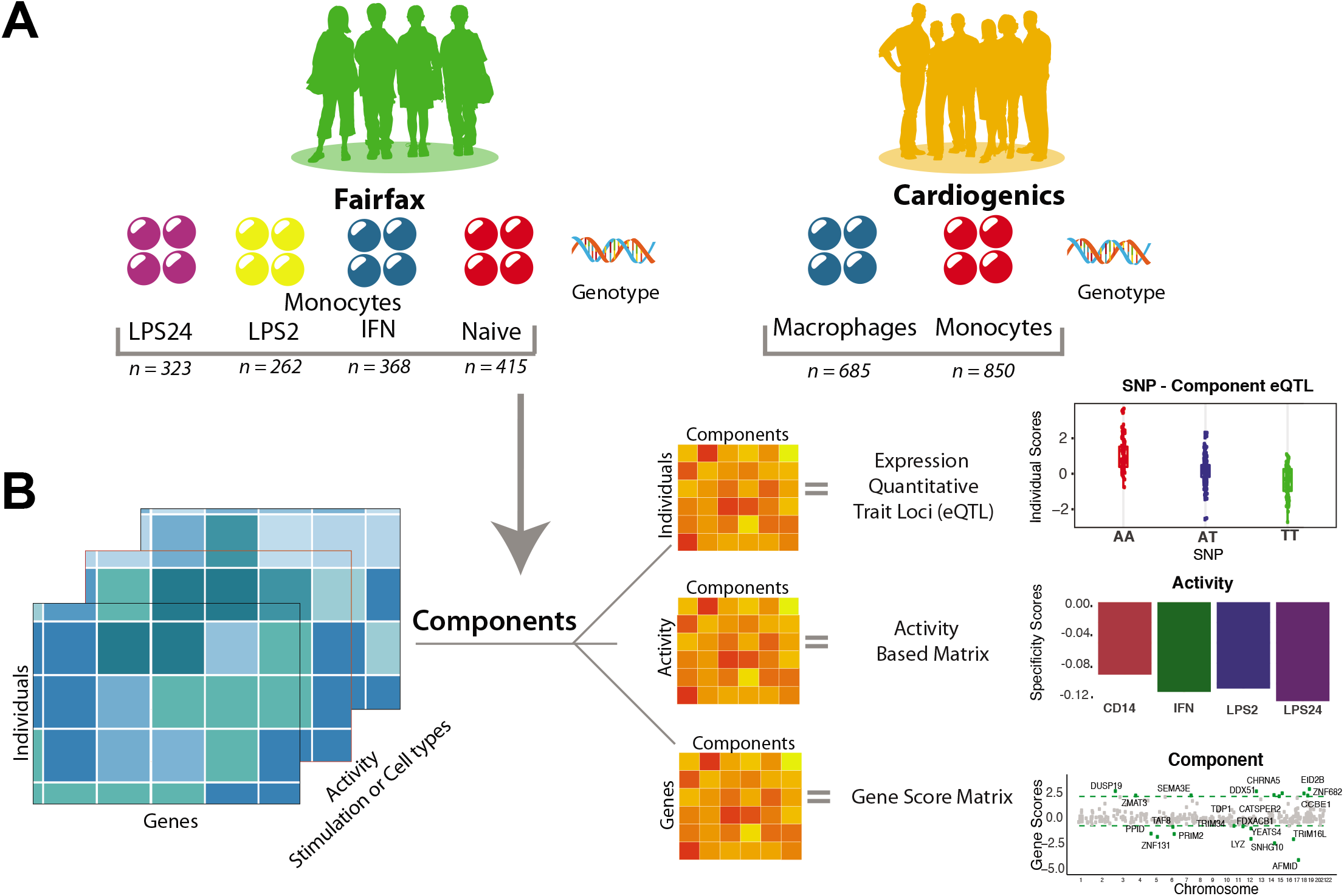
Overview of the study design and method. (A) The gene expression datasets used in this study. Stimulated monocyte gene expression profiles from Fairfax, et al. ^12^ in four conditions: response to lipopolysaccharide at 24 hours (LPS24) and at 2 hours (LPS2), interferon-γ (IFN-γ), and naive (*left panel*). Peripheral blood monocytes (MP) and macrophages (MC) from the Cardiogenics Consortium (*right panel*). (B) Overview of the SDA approach^14^. An illustration of decomposition of gene expression datasets to yield component vectors for relative contribution of each individual, gene and condition. The individual scores matrices are then used as phenotypes with SNP genotypes in order to identify genetic variation correlated with the components (top). The stimulation or cell-activity scores matrix is used to identify the contribution of each condition for the components (middle). The gene scores matrix is used to identify the contribution of each gene within the components (bottom).

We used gene expression profiles from two previously published studies: (1) Fairfax *et al*.^12^ (FF), in which CD14^+^ human monocytes were profiled with two inflammatory stimuli: naïve (CD14), and in response to either interferon-γ (IFN), to lipopolysaccharide at 2 hours (LPS2), or at 24 hours (LPS24); and (2) primary human monocytes and macrophages from the Cardiogenics Consortium (CG)^29^, which includes subjects from a Cardiovascular disease cohort (Fig. 1). The IFN-γ and LPS stimuli were used to elicit an immune reactive phenotype in these cells that may help detect gene networks unidentifiable in the naïve state and better approximate the state of these cells in the context of a degenerating brain. On the other hand, the monocytes and macrophages comparison may help uncover different pathogenic effects of these two cell types. We computed a maximum of 500 components in each dataset, of which the majority were non-sparse (most genes with low or zero gene scores). These non-sparse components capture technical or biological confounders in the expression data (Fig. 2A **and Supplemental Fig. S1**).

**Figure 2:**
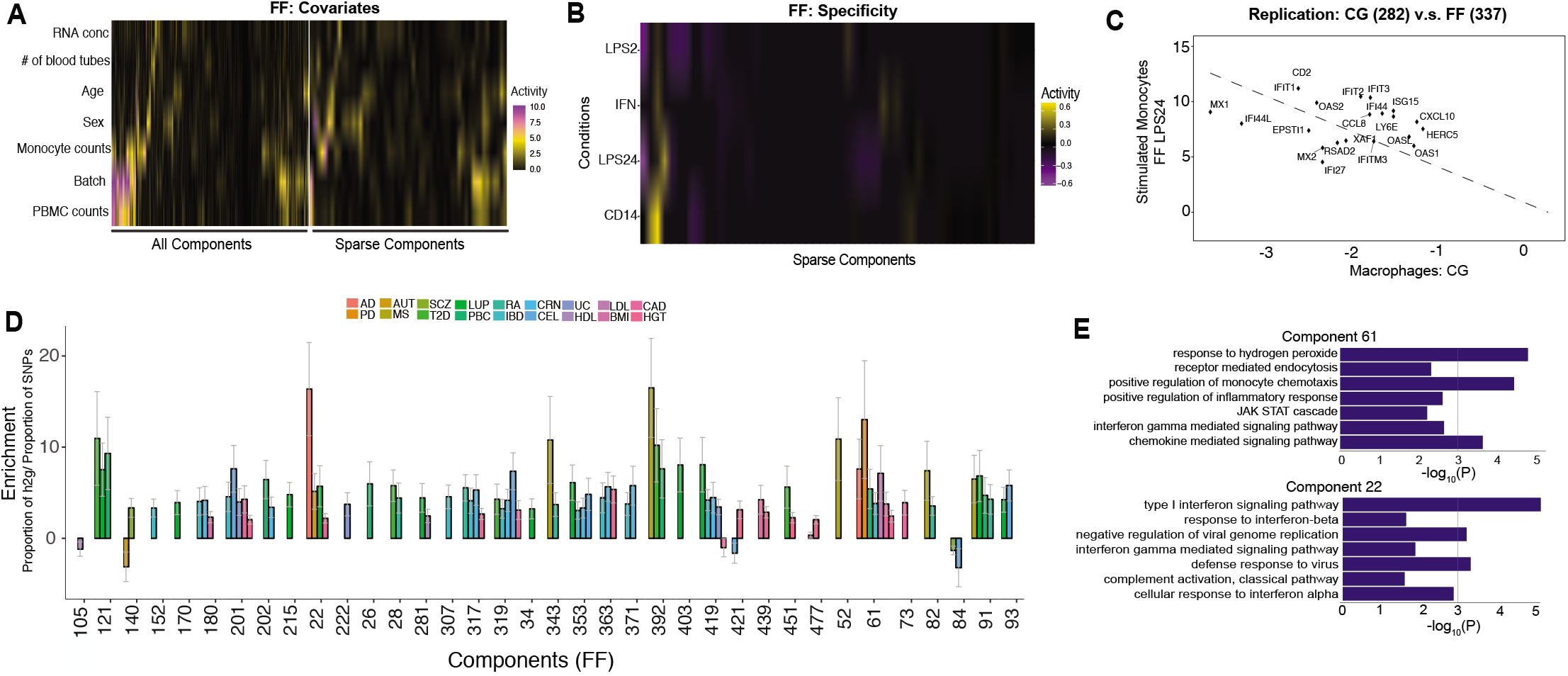
Discovery and reproducibility of sparse components enriched for disease heritability. (A) Heat map of known covariates and correlation with individual scores from each of the 500 components (left) and 56 sparse components (right) in FF data. (B) Heat map of condition specificity scores for the sparse components in FF data. Each row is a stimulation (naïve, LPS, or IFN) and each column is a sparse component. (C) Component replication between component 282 (CG) and component 337 (FF LPS24) for the most highly scored genes. The gene scores from the two components (in CG and FF) are highly correlated. (D) Proportion of heritability for 18 selected complex traits that can be attributed to each sparse component from the FF data. Shown here are enrichment statistics (with standard error) comparing the proportion of SNP heritability within the components divided by the proportion of total SNPs represented at FDR-corrected *P* < 0.05. AD: Alzheimer’s disease; PD: Parkinson’s disease; AUT: Autism; MS: Multiple Sclerosis; SCZ: Schizophrenia; T2D: Type 2 Diabetes; LUP: Lupus; PBC; Primary Biliary Cirrhosis; RA: Rheumatoid Arthritis; IBD: Irritable Bowel Disease; CRN: Crohn’s Disease; CEL: Celiac Disease; UC: Ulcerative Colitis; HDL: High-density lipoprotein Cholesterol; LDL: Low-density lipoprotein Cholesterol; BMI: Body Mass Index; CAD: Coronary Artery Disease; HGT: Height.

Using a ranked-based sparsity statistic (see Methods), we identified 56 and 111 components in FF and CG, respectively, that are sparse, or components with distinguishable non-zero and high scoring genes (**Supplemental Tables S1 and S2**). In the FF dataset we identified 12 sparse components active in naïve, 5-9 active in response to IFN-γ, LPS 2hrs, or LPS 24hrs, and the remaining 25 shared across all conditions (Fig. 2B). In *CG*, 42 sparse components were specific to monocytes and 45 specific to macrophages and 24 were shared between the two cell types (**Supplemental Fig. S1**). Using a two-way reverse correlation approach, we found 7 of the 56 sparse components in FF are conserved between FF and CG (**Supplemental Table S3**). An example of a shared component is shown in Figure 2C.

To assess the functional significance of the components, we tested the genes within each component for enrichment in Gene Ontology (GO) biological processes. We found significant enrichment for GO biological processes for 30 out of 56 sparse components, suggesting that the genes in each sparse component are part of coherent biological processes (**Supplemental Table S4**). To assess the enrichment of disease risk loci in these components, GWAS summary statistics from 18 selected traits and gene-sets from the sparse components were used as input for MAGMA^30^. We found three components significantly enriched for genes in GWAS loci at a Bonferroni-corrected significance threshold and additional four components enriched at a nominal P-value < 0.05 for ten traits (**Supplemental Table S5**). This includes a component with genes in the interferon-signaling pathway, which is enriched for genes in Alzheimer’s disease susceptibility loci. Overall, these results suggest that the sparse components from stimulated monocyte gene expression data have functional significance and, in some cases, may underlie disease-relevant biological processes.

We next reasoned that the monocyte components might contain genes in disease-associated loci that disproportionately contribute to the heritability of complex diseases. We used LD score regression (LDSC)^31^ to partition GWAS heritability into the contribution of SNPs located within genes form each component. We found that 36 out of 56 sparse components in FF showed significant enrichment (FDR-corrected *P* < 0.05) for 18 different traits. This includes a component in FF data (component 61) with significantly enriched for SNP-based heritability for seven traits including Alzheimer’s (7.6-fold, *P* < 0.03), Parkinson’s (13-fold, *P* < 0.01) as well as number of inflammatory and metabolic diseases (Fig. 2D; **Supplemental Table S6**). Another component that is significantly enriched for SNP-based heritability for Alzheimer’s disease (16.4-fold, *P* < 0.04), autism (5.1-fold, *P* < 0.03), and lupus (5.7-fold *P* < 0.03), accounting for 6%, 2% and 2% of SNP-based heritability, respectively. Component 22 is highly enriched for a number of genes in the Type 1 interferon-signaling, defense response to virus, and complement activation pathways (Figure 2E). A number of studies have linked genes from these functional categories to Alzheimer’s disease, autism^32,33^ and lupus^13^. Interestingly, we discovered that this component is also a *trans*-eQTL for Alzheimer’s disease susceptibility loci (see below). Taken together, these results illustrate that several components identified in this study can explain substantial amount of SNP-based genetic heritability for many common diseases and begin to implicate specific innate immune function for common variants across neurodegenerative and inflammatory diseases.

### Detection of context-specific *trans*-eQTLs

To detect *trans*-eQTLs, the individual scores from each component are tested for association with genotype dosages (autosomal SNPs, Minor Allele Frequency [MAF] > 0.01, imputed with the Haplotype Reference Consortium reference panel) across the genome. A stringent quality control (QC) procedure was used to filter out multi-mapped gene probes and non-coding genes prior to *trans*-eQTL analysis (Methods). For clarification, hereinafter, we define the following: a *trans*-eQTL to be an SNP that maps to a component, the relevant SNP itself to be the trans-eSNP and any genes within the component to be trans-eGene(s). In FF, we identified a total of 15,856 *trans*-eQTLs (FDR < 0.05) representing 15,298 unique SNPs (833 independent SNPs; obtained by pruning SNPs in LD with r^2^ < 0.2) and 55 unique sparse components (**Supplemental Table S7**). Of the significant *trans*-eQTLs, approximately 55% are stimuli-specific, detected only in response to either IFN-γ or LPS challenge. For CG, we identified a total of a total of 3,945 *trans*-eQTLs (FDR < 0.05) representing 3,217 unique SNPs (70 independent SNPs; obtained by pruning SNPs in LD with r^2^ < 0.2) and 57 unique sparse components (**Supplementary Table S8**). In each dataset, the components contain anywhere between 5-62 genes with distinguishable non-zero gene scores based on distributional cut-offs (Methods). Our analysis confirmed previously published *trans*-eQTL: a *cis*-eQTL at rs2275888 for *IFNB1* is associated with the expression of 17 genes in *trans* after 24-hour LPS stimulation, many of which are interferon (IFN-β) response genes^2^ (**Supplementary Fig. S2**).

### *Trans*-eQTLs in stimulated monocytes as putative drivers of disease associations

To identify disease risk alleles influencing the expression of the distal gene(s), we investigated whether the *trans*-eSNPs were previously associated with complex traits or diseases. We incorporated 20,094 SNPs from the NHGRI GWAS Catalog pertaining to 1,374 disease or complex traits that were significant at a Bonferroni-corrected significance threshold of *P* < 5 × 10^−8^. We identified 227 *trans*-eQTLs mapping to 29 sparse components that overlap with disease or trait-associated loci (FDR < 0.15; **Supplemental Table S9**). We used a more liberal FDR significance level (0.15) to identify trait-associated *trans*-eQTLs since this class of SNPs has already shown to have functional consequences.

In addition, our FDR correction is applied to testing for genome-wide SNPs rather than just correcting for only the trait-associated SNPs as has been previously done^13^. Nevertheless, to ensure the robustness of the *trans-* eQTLs, we carried out permutation scheme and found 89 *trans*-eQTLs mapping to 18 components were significant at permutation threshold *P* < 1×10^−3^. Fig. 3A provides a subset of the *trans*-eQTLs that are in disease or trait-associated loci. In CG, we found 56 *trans*-eQTLs mapping to 14 sparse components overlapping loci associated with a disease or a complex trait at FDR < 0.15 (**Supplemental Table S10**). Examples of colocalized *trans*-eQTLs with Parkinson’s and Alzheimer’s disease associated susceptibility loci are shown in Figure 3B. We found components that mapped to multiple independent loci and components that mapped to a single locus with multiple disease-associated variants. For example, disease-associated susceptibility alleles in the MHC class II are significantly associated with individual scores of component 363 in FF, which is active in response to all stimuli (**Supplemental Fig. S3**). The component contains several MHC class II genes but also 21 non-MHC genes (e.g., *DEF8* and *AOAH*) with distinguishable non-zero scores. Some of the trans-associations to MHC class II alleles have been previously identified in the SNP-by-gene analysis^12^, but our analysis identified a network of 21 genes pointing to a critical biological pathway underlying disease susceptibility. We identified several other examples of *trans*-eQTLs in disease-associated loci (**Supplemental Figs. S4-S6**). Overall, we were able to not only reproduce previously identified *trans*-eQTLs and individual target genes in monocytes and macrophages but also identify biologically informative gene networks linked to distal genetic variants.

**Figure 3:**
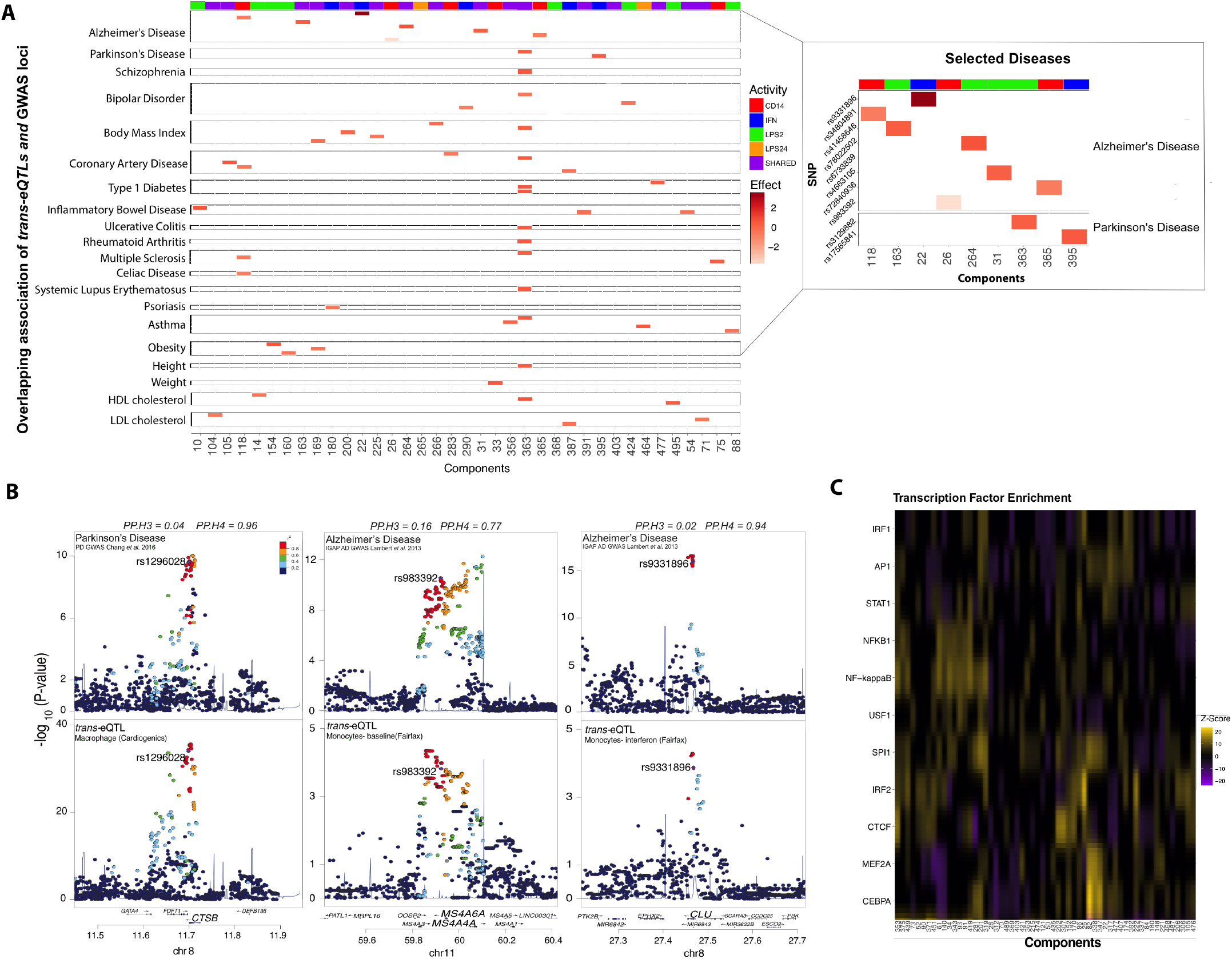
Trans-eQTLs colocalized in disease or trait-associated GWAS loci. (A) Significant *trans*-eQTLs in FF data (FDR < 0.15) in 20 selected disease or trait-associated GWAS loci (*left panel*). Shown are diseases associated trans-eSNP (SNP ID are not shown) on Y-axis and *trans*-eQTL components on X-axis. The red colored boxes reflect the effect size for the *trans*-eQTLs while the horizontal colored header reflects the condition activity scores. *trans*-eQTLs that are in Alzheimer’s or Parkinson’s disease associated loci (*right panel*). Alzheimer’s disease includes GWAS susceptibility loci from Alzheimer’s related traits including the age of onset, age-related cognitive decline, and APOE ε4 carriers. (B) Colocalization of *trans*-eQTLs at Parkinson’s disease susceptibility locus *CTSB* (left panel) and Alzheimer’s disease associated loci *MS4A4A* (middle panel) and *CLU* (right panel). The x-axis in each panel shows the physical position on the chromosome (Mb). The y-axis shows the -log_10_(*P*) association p-values for Parkinson’s disease GWAS from Chang *et al*. 2017 (left panel) and Alzheimer’s disease GWAS from Lambert *et al*. 2013 (middle and right panels). Listed on top are ‘coloc’ posterior probability for hypothesis 3 (PP.H3) and 4 (PP.H4). PP.H3: Association with eQTL and GWAS, two independent casual SNPs. PP.H4: Association with eQTL and GWAS, one shared SNP. (C) Transcription factors whose binding sites occurrence is enriched in the target set of genes within each sparse component compared to the expected occurrence estimated from a background set.

Of the disease-associated *trans*-eSNPs (223 in FF and 43 in CG), 85 and 27 in FF and CG, respectively, were also *cis*-eSNPs for nearby genes. We applied a Mendelian randomization method^34^ to quantify support for a direct relationship between each SNP that has a *cis*-eQTL to their trans-eGene(s). We applied Mendelian randomization to test if the trans-eSNP mediated the effect on the trans-eGene(s) through their respective *cis*-gene using two different approaches. First, we used the individual scores from SDA for the 56 sparse components in the mediation analysis. At a nominal *P* < 0.05, we found 35 and 15 in FF and CG, respectively, where the *cis*-gene is a significant mediator of the component individual scores. In the second analysis, we directly tested for mediation using the expression level of each *trans-eGene* within each respective component. From this directed approach, we found a significant mediation effect for at least one *trans-eGene* for 79 and 21 in FF and CG trans-eSNPs, respectively. Of the significant *trans* mediators, we found that only 11 genes were transcription factors (TF), suggesting that TFs are not primary regulators of the *trans* network. However, we observed regulatory motifs for the same TF enriched among the all trans-eGenes within each network (Fig. 3C). These include many myeloid lineage-specific factors (e.g., *PU.1, C/EBPα and β*, and *MEF2A*) and interferon-related (e.g., *IRF1, IRF2*, and *STAT1*) factors (Fig. 3C). In summary, our results are consistent with other findings that TFs are not the primary driver of *trans*-eQTL networks but are enriched for specific TF binding motifs, suggesting that the trans-eGenes are under the same regulatory control.

### Macrophage-specific *Trans*-eQTL at Parkinson’s disease susceptibility loci

Considering that GWAS have identified loci with genes related to the innate immune system both in Alzheimer’s disease and Parkinson’s disease, we hypothesize that analysis of monocytes and macrophages could unravel new gene-sets associated with disease susceptibility. In support of this hypothesis, we observed in CG macrophages that the rs1296028 affects the expression of the lysosomal protease Cathepsin B (*CTSB*) (*P* = 1. 46 x 10^−22^; FDR = 1.31×10^−18^) in *cis* (**Supplemental Fig. S7**) and the expression of 16 genes in *trans* (*P* = 8.29 x 10^−35^; FDR = 9.48×10^−28^) (Figs. 4A and 4B). The lead trans-eSNP rs1296028 (MAF = 0.11) at the *CTSB* locus is associated with Parkinson’s disease susceptibility (in LD with GWAS lead SNP rs2740594; r^2^ = 0.68)^22^. Both *trans*-eQTL and disease association signal were consistent with a model for shared causal variants (as indicated by coloc^35^ posterior probability PP3+PP4 = 0.99, PP4/PP3 = 24) (Fig. 3B). Although we did not find significant enrichment of any specific biological pathways with the Ingenuity Pathway Analysis (IPA) tool, we found enrichment for biological processes such as ‘*lysosomal pathway*’ (*P* = 5.2×10^−3^) and ‘*cholesterol degradation*’ (*P* = 6.22×10^−4^) when we lowered the threshold for gene scores. The directions of effect for all the genes in this network were consistent: the minor allele (rs1296028-G) is associated with decreased expression of all 16 genes (Fig. 4A; **Supplemental Figs. S8 and S9**). Using a small independent macrophage eQTL dataset (STARNET^5^; n=82), we were able to replicate 4 of the 16 *trans*-eQTL associations (FDR < 0.20; **Supplementary Fig. S10**). The direction of effect is consistent across the two datasets: rs1296028-G is associated with increased expression of the target genes as well as decreased risk for PD.

**Figure 4.**
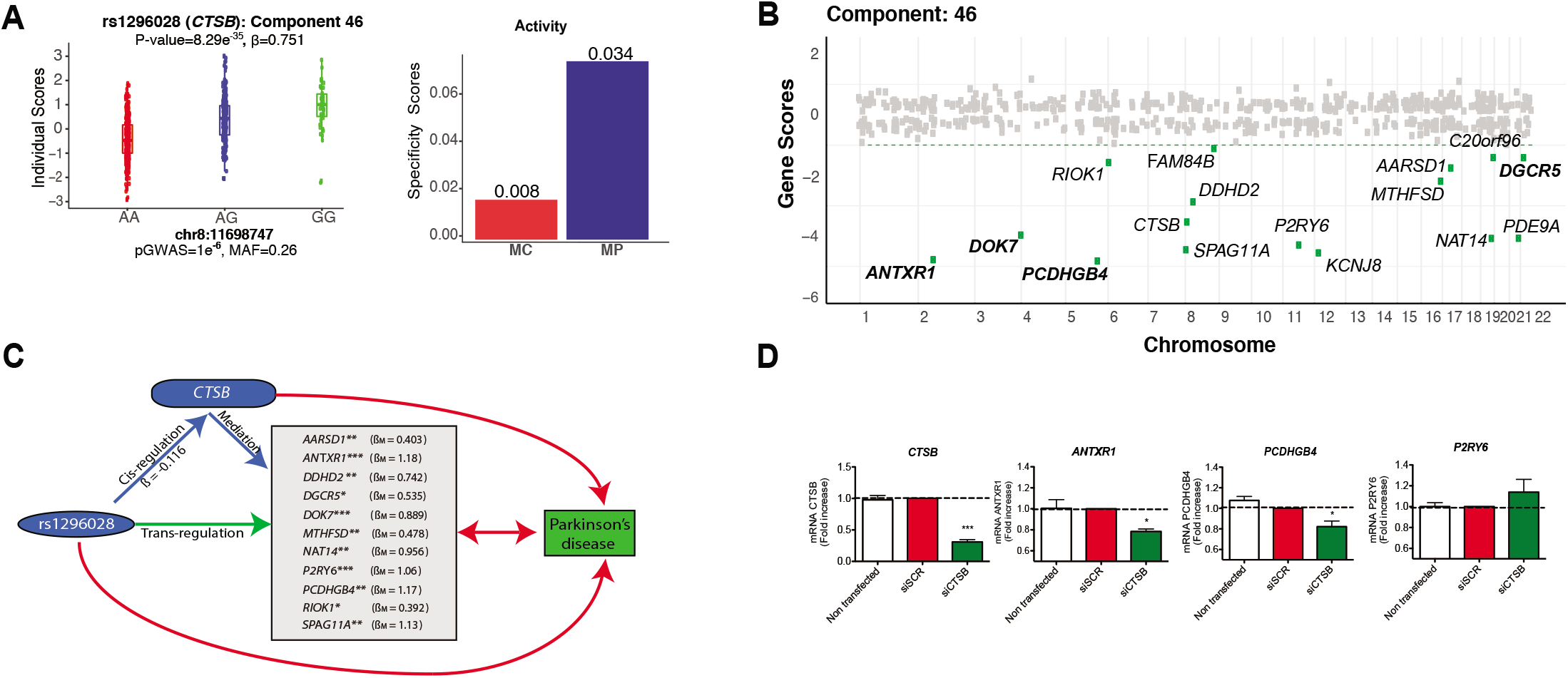
Macrophage-specific *trans*-eQTL colocalized in Parkinson’s disease associated *CTSB* locus. (A) The genotype for Parkinson’s disease susceptibility allele rs1296028 is significantly associated with the individual scores of CG component 46. The rs1296028 affects the expression of 16 genes in component 46 in macrophage (CG) (left panel). Tissue specificity scores suggest CG component 46 is active in macrophage (Right). *P*-value; ***: <1e^−50^ I **: <1e^−10^ I *: <1e^−5^ (B) Gene scores for component 46 across the genome. Only the *trans-eGenes* with PIP > 0.5 and 2.5% distributional cut-off (green dotted line) are shown. *Trans-eQTL* associations that replicated (FDR < 0.20) in an independent STARNET macrophage dataset are shown in bold (C) Parkinson’s disease SNP rs1296028 mediates *trans-effects* to 11 genes through cis-mediator cathepsin B (*CTSB*). The beta coefficients from Mendelian randomization analysis are shown for the significant *trans-eGenes*. (D) Experimental validation using THP-1 derived macrophages. *CTSB* was knock-down using siRNA during 48 h and the levels of the top-scoring genes in the component were measured by qPCR. Data was normalized against scramble siRNA (SCR). *P*-value; *: <0.05 I ***: <0.001.

Using Mendelian randomization, we found mediation effect for rs1296028 through *CTSB* for 11 of the genes in this network, of which the most significant mediation effects were for *ANTXR1, PCDHGB4, SPAG11A* and *P2RY6* (Fig. 4C). To examine the causality of this relationship between *CTSB* and the *trans* target genes, we performed siRNA-mediated knockdown of *CTSB* and measured the effect on the four previous mentioned genes that have β_M_ > 1 using human THP-1 derived macrophages. We observed a concomitant reduction in *ANTXR1* and *PCDHGB4* mRNA levels (*P* < 0.05) as measured by qRT-PCR (Fig. 4D). A non-significant effect was observed in *P2RY6*, whereas *SPAG11A* is not expressed in the THP-1 cell line. The knockdown experiment provides an independent validation for some of the genes in the *CTSB *trans*-eQTL* network. However, additional experimental validation is necessary to fully elucidate the role of *CTSB* lysosomal network in the etiology of Parkinson’s disease.

### Alzheimer’s disease loci affect the expression of genes in Type 1 interferon signaling pathway

Following the same rationale, we attempted to find altered monocyte and macrophage gene networks associated with Alzheimer’s disease susceptibility alleles. We found eight *trans*-eQTLs that overlap with Alzheimer’s disease (or Alzheimer’s disease-related phenotypes) susceptibility loci (Fig. 3A), and we highlight two examples (Figs. 5 and 6). First, we observed that rs983392 on chromosome 11q12, reported to be associated with Alzheimer’s disease (*MAF* = 0.41, *P*_gwas_ = 6×10^−16^), is a *trans*-eQTL to a component with 54 genes of distinguishable non-zero loading scores (*P* = 1.5×10^−4^; FDR < 0.15) (Figs. 5A and 5B).

**Figure 5.**
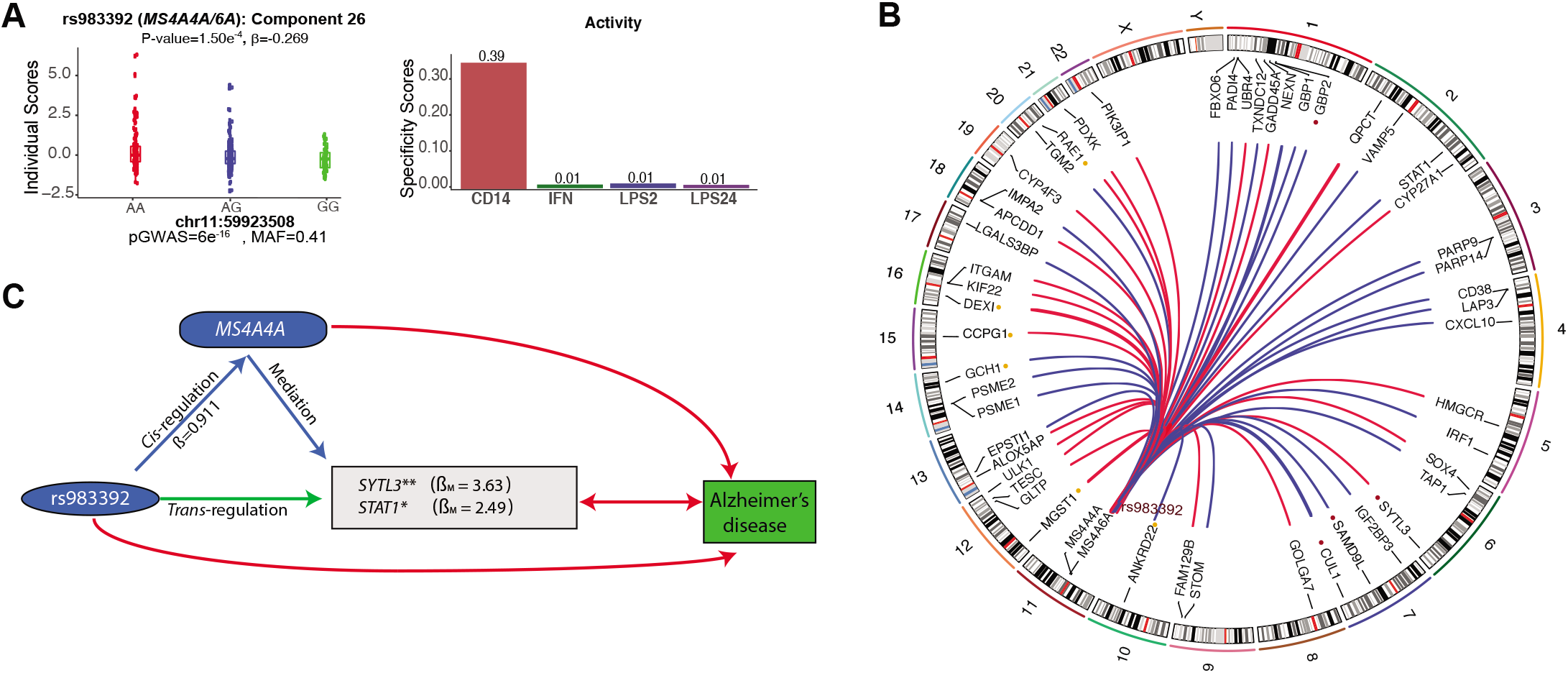
*trans*-eQTL colocalized in Alzheimer’s disease associated *MS4A* locus. (A) The genotypes for Alzheimer’s disease susceptibility allele rs983392 are significantly associated with the individual scores of component 26. rs983392 is *trans*-eQTL to FF component 26 with 54 genes (*left panel*). The component 26 is active only at baseline (*right panel*). (B) Circular plot demonstrating the chromosomal position of trans-eSNP (rs983392) and the 54 *trans* target genes. The minor and AD-protective allele rs983392-G is associated with decreased (blue lines) and increased (red lines) expression of *trans*-eGenes. The colored dots are *trans*-eQTL that replicates in ImmVar baseline monocytes (red and yellow color dots denote *trans*-eQTL association at FDR < 0.05 and 0.20, respectively). Only the *trans*-eGenes with PIP > 0.5 and 2.5% distributional cut-off are shown. (C) Alzheimer’s disease SNP rs983392 mediates trans-effects to two trans-eGenes through cis-mediator *MS4A4A*.

**Figure 6.**
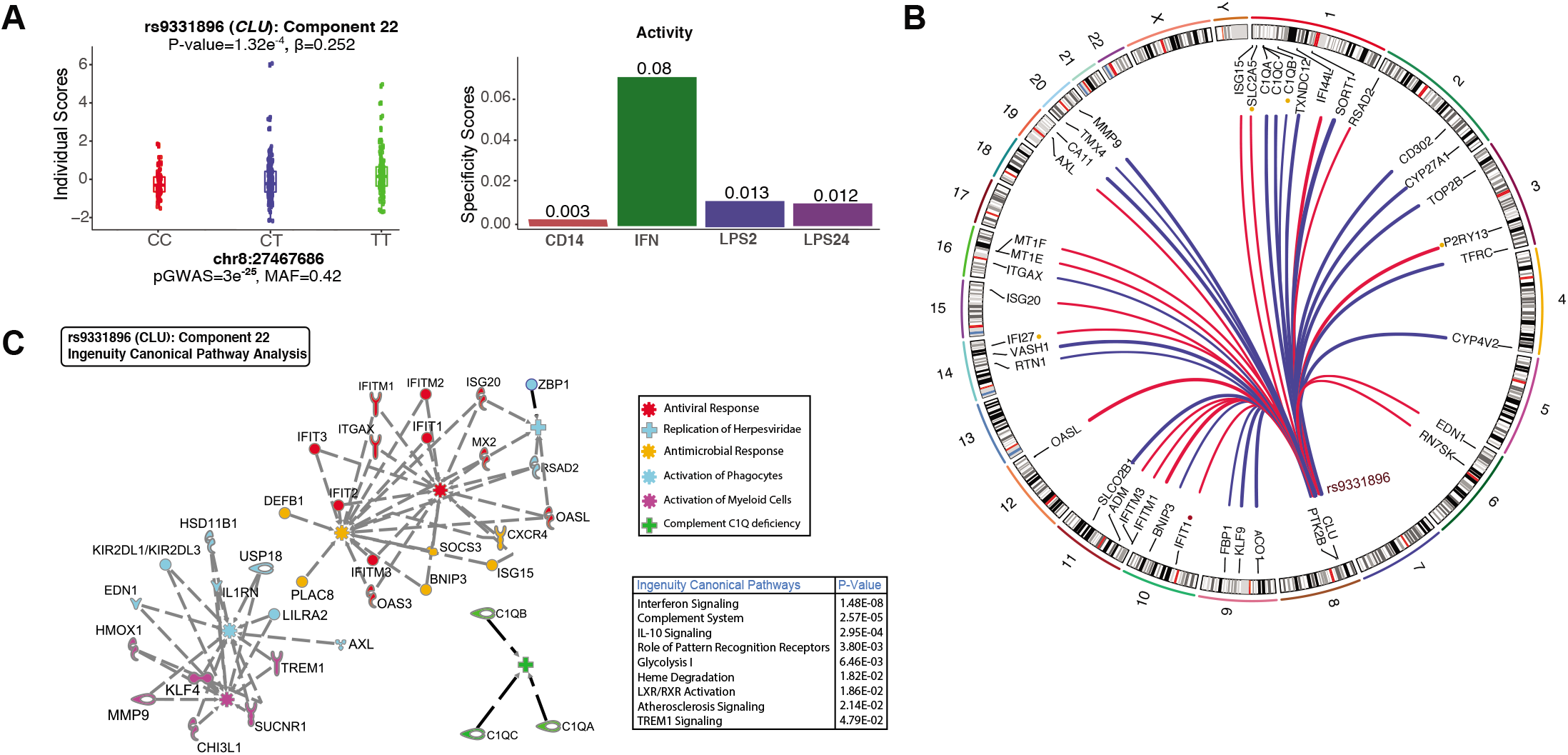
Trans-eQTL colocalized in Alzheimer’s disease associated *CLU* locus. (A) The genotypes for Alzheimer’s disease susceptibility allele rs9331896 are significantly associated with the individual scores of component 22 (*left panel*). The component is active in monocytes (FF) in response to interferon-γ (*right panel*). (B). Circular plot demonstrating the chromosomal position of ŕrans-eSNP (rs9331896) and the 38 *trans* target genes. The minor and AD-protective allele rs9331896-C is associated with decreased (blue lines) and increased (red lines) expression of *trans*-eGenes. The colored dots are *trans*-eQTL that replicates in ImmVar IFN stimulated monocytes (red and yellow color dots denote *trans*-eQTL association at FDR < 0.05 and 0.20, respectively). Only the *trans-eGenes* with PIP > 0.5 and 2.5% distributional cut-off are shown. (C) IPA pathway analysis of the *trans-eGenes* in this component. The IPA canonical pathway enrichment P-values are shown in the lower right. The pathway network is grouped by IPA biological functions. The functional groups are defined by different colors and symbols (top right).

The component is active in the naïve condition, and the stimulation of IFN-γ or LPS ablates the *trans*-eQTL signal. A significant *cis*-eQTL effect for rs983392 to both *MS4A4A* (FDR = 2×10^−3^, β = -4.4) and *MS4A6A* (FDR = 5×10^−4^, β = -4.7) is observed only in the naïve condition (**Supplemental Fig. S11**). Both *trans*-eQTL and disease association signal were consistent with a model for shared causal variants (as indicated by coloc^35^ posterior probability PP3+PP4 = 0.93, PP4/PP3 = 4.8) (Fig. 3B). The top-scoring genes in this component encode for Interferon-Inducible Guanylate Binding Proteins (*GBP1, GBP2, GBP4*, and *GBP5*), followed by *STAT1* and *IRF1*, which are key TFs for IFN-γ activation and type 1 interferon signaling (Fig. 5B **and Supplemental Figs. S12-S14**). The genes in this component are enriched for IPA biological processes such as ‘*interferon signaling*’ (*P* = 5.77×10’^16^) and the ‘*antigen presentation*’ pathway (*P* = 6.62×10’^9^). IPA also identified enrichment for several annotated biological functions including ‘*antiviral response*’ (*P* = 1.26×10^−33^). To determine if the observed *trans*-eQTL associations are mediated by expression of either *MS4A4A* or *MS4A6A* in *cis*, we performed Mendelian randomization-based mediation for using both component individual scores and gene expression levels. We observed trans-regulatory effects of rs983392 on *STAT1* and *SYTL3* mediated through *MS4A4A* but not through *MS4A6A* (Fig. 5C).

Another Alzheimer’s disease-associated variant rs9331896-C (MAF=0.41, *P*_gwas_ = 3×10^−25^) on located on chromosome 8p21 within intron 2 of the *CLU* gene is associated with the individual scores of a component consisting of 38 genes with non-zero scores (*P* = 1.32×10^−4^; FDR < 0.15) (Figs. 6A and 6B). This component is active in IFN-γ stimulated monocytes (Fig. 6B). Both *trans*-eQTL and disease association signal were consistent with a model for shared causal variants (as indicated by coloc^35^ posterior probability PP3+PP4=0.96, PP4/PP3 = 47) (Fig. 3B). The SNP rs9331896-C had no *cis*-eQTL effect on *CLU* in monocytes (or in CG macrophage) but we have reported previously a splicing QTL for *CLU* in *DLPFC^36^* and more recently we observed the same splicing event in human microglia (unpublished). Given these results, it is likely that the *cis* effect that we observed could mediate the trans-effect through the expression of specific isoform. The top-scoring genes in the network include the members of Interferon Induced Transmembrane Proteins (*IFITM1* and *IFITM3*) followed by *MT1E, MT1X*, and *MT1G and OASL*, all essential proteins involved in the innate immune response to viral infection (Fig. 6B; **Supplemental Figs. S15-S17**). Using IPA we found significant enrichment for ‘*interferon signaling*’ (*P* < 1.48×10^−8^) and ‘*complement cascade*’ pathways (*P* < 2.57×10^−5^) (Fig. 6C). This component also contained genes in the complement components C1q family of genes (*C1QA, C1QB*, and *C1QC*). Interestingly, protein products of *CLU* (previously known as CLI for complement lysis inhibitor) and complement component genes physically interact and are part of a protein-protein interaction (PPI) network (**Supplemental Fig. S18**), providing an independent validation of our *trans*-eQTL effects. Finally, we observed several genes in this component enriched for IPA biological processes such as ‘*Activation of Phagocytes*’ including *AXL* and *TREM1* (Fig. 6C).

To assess the robustness of the *trans*-eQTLs that colocalize with Alzheimer’s disease risk alleles, we attempted to replicate the association signals using an independent dataset from the ImmVar study ^7^. Of the 54 trans-eGenes in the *MS4A4A/6A* component, the expression of 10 trans-eGenes in naive monocytes were significantly associated with rs983392 (FDR < 0.20; **Supplemental Fig. S19**). Many of the *trans*-eQTLs were highly significant in the ImmVar data including *GBP2, CUL1*, and *SYTL3* (Fig. 5B). Of the 38 trans-eGenes in the *CLU* component, expression levels of five genes were significantly associated with rs9331896 at FDR < 0.20 in the IFN stimulated monocytes (**Supplemental Fig. S20**). Despite the small size and differences in stimuli in the replication dataset, we were able to detect suggestive association signal for a small number of *trans* effects.

## Discussion

Here, we used tensor decomposition methodology^14^ to detect *trans*-eQTL in naïve and stimulated human monocytes and macrophages. We detected hundreds of *trans*-acting regulatory effects on a number of genes in sparse components. These components explain substantial amount of SNP-based genetic heritability for many common diseases. After examining the genes in each component, we observed that the target genes were involved with coherent biological processes and had regulatory motifs that are enriched for the same transcription factor. The majority of the *trans*-eQTLs were hotspot loci, each of which altered the expression of many genes within our sparse components. We detected significantly more *trans*-eQTLs in stimulated monocytes compared to naïve monocytes or macrophages despite twice the sample size for CG. Among the significant *trans*-eQTLs, we found 55% were stimuli-specific, suggesting that a larger number of *trans*-eQTLs are detectable only in the presence of specific stimuli. These observations are consistent with findings from other species such as *Caenorhabditis elegans^37^* and *Saccharomyces cerevisiae^38^* showing that environmental perturbation yields a higher number of *trans*-eQTLs compared to cis-eQTLs. Together, these results underscore the need to perturb primary cells with environmental stimuli to discover genotype-phenotype relationships in *trans*.

The target trans-eGenes reveal biological processes and downstream effects for a number of disease-associated susceptibility alleles. This includes Parkinson’s disease associated *trans*-eQTLs with 16 target genes mediated by the lysosomal protease Cathepsin B (*CTSB*). We corroborate a previously reported *cis-* eQTL effect on *CTSB* driven by a Parkinson’s disease-associated genetic variant^22,28^. However, the main contribution of this study is our ability to detect reproducible Parkinson’s disease associated *trans*-eQTLs and experimentally validate several of the target genes. Nevertheless, we are still unable to fully understand this gene network in the context of the disease due to our incomplete understanding of the functions of many genes within the network. The only gene in this network known have any functional interaction with *CTSB* to the date is *ANTXR2* (same family of *ANTXR1*, found in the network), where *CTSB* -mediated autophagy flux facilitates the delivery of toxins into the cytoplasm^39^. Further functional studies will be necessary to validate not only the *trans* targets but also to understand mechanisms underlying the Parkinson’s disease susceptibility at the *CTSB* locus. While alterations in autophagy and lysosomal pathways have been widely reported in neurons from Parkinson’s disease^40^, our results open future mechanistic studies focusing on macrophage function and gene networks in Parkinson’s disease.

We have previously reported that a number of Alzheimer’s disease-associated susceptibility alleles colocalize with cis-eQTLs in peripheral monocytes^7,41–43^. Here we hypothesized that Alzheimer’s disease risk and protective alleles may modulate myeloid cell function within specific biological pathways. In support of this hypothesis, our analysis showed that the Alzheimer’s disease susceptibility alleles at the *MS4A4A/6A* and *CLU* loci are associated with the individual scores of two sparse components. The component containing type 1 interferon genes explains substantial proportion (6%) of SNP-based genetic heritability for Alzheimer’s disease. These *trans* target genes for both loci intersect with interferon-related functional genes that are responders of IFN-γ, key regulators of the IFN response (*IRF1* and *STAT1*), and type I interferon and antiviral effectors (OAS, *IFIT*, and *GBP* families). These findings confirm previous results in the p25/Cdk5 model of neurodegeneration where a reactive microglia phenotype with activated IFN pathway was found^44^. Given that altered IFN signaling has been detected in both modules, our results suggest that dysregulation of this pathway in microglia might have a role in AD pathogenesis.

The Alzheimer’s disease-associated *trans*-eQTL in the clusterin (*CLU/APOJ*) locus is also associated with the expression of genes of the complement cascade. The role of the complement system in Alzheimer’s disease has been suggested mostly by mouse studies in which microglia are thought to be the cellular effector of complement-mediated synaptic loss in Alzheimer’s disease ^45^. Indeed, C1q and oligomeric forms of amyloid-β operate in a common pathway to activate the complement cascade and drive synapse elimination by microglia^45^. In addition, a post-mortem study of Alzheimer’s disease brains showed increased expression levels for complement components in the Alzheimer’s disease brains^46^. Other genes in this component include marker genes (AXL and *ITGAX*) for damage-associated microglia, or DAM, a recently identified subset of microglia found at sites of neurodegeneration^24^. Both *AXL* and *ITGAX* are key genes involved in the Trem2-dependent DAM program, which involves upregulation of phagocytic and lipid metabolism genes^24^. The genes in the clusterin mediated network may have a role in debris clearance such as the removal of apoptotic neurons or Aβ aggregates by microglia. Further work is necessary to fully understand the role of complement and interferon signaling genes in the development and progression of AD pathology.

With this study, we demonstrate that gene expression from peripheral immune cells are a valuable source to study gene networks associated to different diseases, including neurodegenerative diseases. Nevertheless, this study has several limitations. First, while we were able to detect robust gene networks linked to distal genetic variants, we are still underpowered to detect *trans* target genes with smaller effect sizes. Our results suggest that many more genes have non-zero gene scores and are likely to contribute to the *trans*-eQTL networks but we are unable to detect them with the current sample size. We estimate that thousands of individuals would be needed to reliably detect effect sizes that explain a small proportion of the *trans*-QTL variance. Secondly, the reproducibility of the *trans*-eQTLs is still challenging without a stimulated dataset of comparable sample size. Thus, as larger datasets become available it will be important to validate our catalog of *trans*-eQTLs. Third, it is not clear if the *trans*-eQTLs networks identified in peripheral monocytes will be conserved in microglia during the transition to a reactive state under conditions of brain-tissue damage encountered during aging or neurodegeneration. These networks in monocytes may play a direct role in the pathogenesis of Alzheimer’s or Parkinson’s disease or, given the shared ontogeny of these two cell lineages, may serve as a proxy for microglial activities within the healthy, aged or diseased brain. While many genes are expressed in both peripheral monocytes and macrophages and CNS microglia, some of the genes are markers for microglia in the healthy brain, or are active during the transition from the homeostasis-associated state to a brain damage-response state. Future studies incorporating transcriptome profiles from primary human microglia from autopsied samples, or Induced Pluripotent Stem Cells (iPSC)-derived microglia challenged with disease-relevant stimuli will be an important resource in uncovering *trans*-eQTL networks underlying neurodegenerative diseases.

In summary, we identified robust *trans*-eQTL networks in peripheral myeloid cells that reveal downstream biological processes of several disease-associated loci. Although further mechanistic work is necessary to validate these gene networks our findings provide compelling human genetic evidence for lysosomal pathway contributing to Parkinson’s disease, and for myeloid phagocytosis, complement cascade and type I interferon-mediated signaling pathways contributing to Alzheimer’s disease.

## Methods

### Overview of Sparse Tensor Decomposition

We applied a sparse tensor decomposition model^14^ to deconstruct multiway gene expression data into latent components or *objects* of smaller dimension for simultaneous analysis. The model itself is 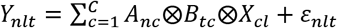, where C is the number of components and A is an *N×C* matrix of individual scores, B is a *T×C* matrix of tissue scores and X is a *C×L* matrix of gene loadings. The error term is modeled as 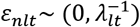, where *λ_lt_* is the precision of the error term at the *l*^th^ gene in the *t*^th^ tissue. An indicator variable *I_nt_* that equals 1 when gene expression has been measured in tissue *t* for sample *n* and 0 otherwise. The likelihood is then given by 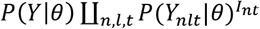, where *θ* is the vector of model parameters. This model is fit in a Bayesian framework, and place priors on the entries of the matrices *A, B, X* and also the precisions *λ_lt_*. A key prior is the one placed on the elements of the gene loadings matrix *X*. A hierarchical ‘*spike and slab*’ prior^28^ is used to encourage sparsity in the rows of matrix *X*. The ‘*spike and slab*’ prior allows to shrink gene loadings to zero to infer more clearly which genes are involved in each component. See Hore *et al*. ^14^ for further details of the model.

We used the Sparse Decomposition of Arrays (SDA) software package (see URLs) to deconstruct multi-way gene expression data into latent components. The SDA model allows for non-sparse components (genes with close to ‘0’ loading scores) that might arise as a result of confounding effects, such as batch effects or technical artifacts. To exclude the non-sparse components from our *trans*-eQTL analysis, we implemented a sparsity ranking statistic alongside the posterior inclusion probability (PIP) > 0.5 inclusion component probability and 2.5% distributional cut-off for gene scores. The statistic is 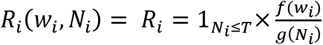 *for i* = 1,…, *n*; where *w_i_* is a weight function, *N_i_* are the number of (non-zero) genes. A cutoff value is derived by simply finding a lower bound through a limiting case instead of using distributional assumptions. See Supplemental Notes 1 for further details.

### Gene Expression Data

#### Fairfax

We obtained processed microarray gene expression data from the Fairfax et al. ^12^ from ArrayExpress (E-MTAB-2232). In the Fairfax dataset, the gene expression of primary human monocytes was profiled in four conditions: naive, in response to interferon-γ, and to lipopolysaccharide at 2 hours, and at 24 hours. Of the 432 total, gene expression profiles were available for 414, 367, 261 and 322 for baseline, IFN-γ, LPS2hr and LPS 24hr, respectively. The gene expression data was generated using Illumina HumanHT-12 v4 BeadChip gene expression array platform with 47,231 probes. Of these, 28,688 probes correspond to coding *transcripts* with well-established annotations and map unequivocally to one single genomic position were kept. We obtained the GRCh37 start and end coordinates for those genes from Ensembl for eQTL analysis. We kept the maximum of median probe expression (across individuals) for multiple probes (mapping to the same gene). This resulted in 17,509 genes used as an input analysis for SDA. See Fairfax, et al. **^12^** for further details on microarray data quality control.

#### Cardiogenics

Cardiogenics is an European collaborative project, it started in January 2007 and was funded by the European Commission through its Sixth Framework Program (reference LSHM -CT-2006-037593). As part of the Cardiogenics project, RNA from monocytes and macrophages of patients with coronary artery disease and healthy individuals was prepared and genome-wide expression was assessed in both cell types using the Illumina HumanRef 8 v3 Beadchip containing 24,516 probes corresponding to 18,311 distinct genes and 21,793 Ref Seq annotated transcripts. The DNA of all these individuals was genotyped using the Human 610 Quad custom arrays. We obtained processed microarray gene expression data from the European Genome-phenome Archive (Study: EGAS00001000411; Dataset ID: EGAD00010000446, EGAD00010000448 and EGAD00010000450). The details of the Cardiogenics datasets can be found in Rotival *et al.*^16^ and Garnier *et al*^29^.

#### ImmVar

The Immune Variation (ImmVar) project consists of 162 African American subjects of European and African ancestry, 155 East Asian subjects of Chinese, Japanese or Korean ancestry, and 377 Caucasian subjects of European ancestry. Genome-wide genotyping was done using Illumina Infinium Human OmniExpress Exome BeadChip and subsequently imputed using the Michigan Imputation Service with Human Reference Consortium v1.1 reference panel. The mRNAs were profiled by Affymetrix GeneChip Human Gene ST 1.0 microarrays and raw data CEL files were processed using the Robust Multichip Average algorithm in Affymetric PowerTools. For further details see Raj, et al. ^7^. In addition to array data, the stimulated and baseline monocytes from the ImmVar cohort were subsequently sequenced^47^. The RNA-seq counts of baseline and interferon stimulated ImmVar data were downloaded from accession GSE92904. The raw fastq and genotype data is available from dbGAP under accession phs000815.v1.p1.

#### STARNET

The Stockholm-Tartu Atherosclerosis Reverse Network Engineering Task (STARNET)^5^ macrophage gene expression data and genotype data is available from dbGAP under accession phs001203.v1.p1.

#### Genotype Quality Control and Imputation

The raw genotype data were downloaded from the EGA (EGAS00000000109 and EGAD00010000450) for Fairfax and Cardiogenics, respectively, and from dbGAP (accession phs001203.v1.p1) for STARNET. The raw data was subsequently imputed using the Michigan Imputation Service with Human Reference Consortium v1.1 reference panel of European ancestry. We used 5,386,706 variants with minor allele frequency >= 0.01, INFO score >= 0.3 and Hardy-Weinberg equilibrium chi-sgưare *P* > 1×10^−6^ for downstream eQTL analysis.

#### Gene Expression Quality Control

The gene expression array data sets were processed using the same pipeline. We performed an additional level of probe set filtering: (i) all array probes with a single nucleotide polymorphism (SNP) at minor allele frequency (MAF) greater than 0.1 in any of the 1000 Genomes populations were removed. (ii) probes that do not map to the human genome were removed. (iii) potential cross-hybridization probes, as provided by Affymetrix or Illumina, were flagged and removed prior to the *trans*-eQTL association analysis. (iv) only uniquely mapping probes and those mapping to GENECODE v19 were included. (v) probes mapping to the X and Y chromosome were excluded. The gene expression data from each study were first internally normalized by dividing the expression values for each gene in individuals of that cohort by the mean expression value across the study, with the assumption that inter-batch differences on normalized data are much lower than those on raw expression values. These normalized values for the three cohorts were assembled and log_2_-transformed.

#### GWAS SNPs

The list of SNPs associated with various human complex diseases and traits was downloaded from the GWAS Catalog (see URLs; accessed October 2017). We included the SNPs with genome-wide significant association (*P* < 5×10^−8^) in our analyses. The list of SNPs was pruned to eliminate SNPs with high LD (pair-wise r^2^ < 0.4).

#### eQTL Mapping

Prior to testing for *trans*-eQTLs association, we discarded the components that were correlated with known biological or technical covariates (see Figure 1). Only SNPs with a minor allele frequency (MAF) > 0.05 and a Hardy-Weinberg equilibrium *P* > 0.001 were included in the analyses. We performed association tests of SNP genotype or imputed allele dosage with individual scores using linear regression as implemented in the Matrix-eQTL^16^ software package. For genome-wide *trans* analyses, we used an FDR of 0.05 to report the significant associations. We used a more liberal FDR threshold of 0.15 for *trans*-eQTLs that colocalize with disease-associated loci. For *trans*-eQTLs that colocalized in disease-associated loci, we permuted individual scores (from SDA) 10,000 times to generate a null-distribution. We then compared the nominal eQTL *P*-values to the empirical distribution created from the permuted datasets^8^.

The *cis* and *trans* eQTL (SNP-by-gene) was carried using a linear regression to perform associations between the imputed SNPs and the normalized gene expression. *cis*-eQTL analysis was limited to SNPs located within 1MB either side of the transcription start or end site. We used PEER^48^ to account for confounding factors in the gene expression data. We fit the PEER model to gene expression data with 15 factors. The residuals from PEER-corrected gene expression data and imputed SNP dosages were used to perform linear regression using the Matrix-eQTL^49^ software package. FDR for the *cis-* and *trans*-eQTL analysis was calculated following Benjamini and Hochberg^28^ procedure as implemented in the Matrix-eQTL package.

#### Enrichment Analysis

We used oPOSSUM-3^50^ to identify over-represented transcription factor binding sites (TFBS) among genes in each component. The Z-score uses the normal approximation to the binomial distribution to compare the rate of occurrence of a TFBS in the target set of genes (in each component) to the expected rate estimated from the precomputed background set. TopGO package^51^ was used to perform enrichment analysis for Gene Ontology (GO) terms. We ran enrichment for significant (PIP > 0.5 and distributional cut-off 2.5%) genes amongst all component networks. We also applied Ingenuity Canonical Pathway (IPA) analysis to perform pathway enrichment for genes within each component. We used MAGMA^30^ to test for enrichment of components among GWAS traits.

#### LD Score Regression

We used stratified LD score regression^31,52^ to partition SNP-based disease heritability within categories defined by the sparse components. Using GWAS summary statistics from 18 traits or complex diseases (Alzheimer’s disease, Parkinson’s disease, Autism, Multiple Sclerosis, Schizophrenia, Type 2 Diabetes, Lupus, Primary Biliary Cirrhosis, Rheumatoid Arthritis, Irritable Bowel Disease, Crohn’s disease, Celiac Disease, Ulcerative Colitis, High-density lipoprotein Cholesterol, Low-density lipoprotein Cholesterol, Body Mass Index, Coronary Artery Disease, and Height) and LD modeled from 1000 genomes reference panel of European ancestry, we calculated the proportion of genome-wide SNP-based heritability that be attributed to SNPs within each component. Categories for each component were defined by taking all the SNPs (within each gene plus 10 kb +/- from transcript start and stop sites) for all genes within the component. To improve model accuracy, the categories defined by components categories were added to the ‘full baseline model’ which included 53 functional categories capturing a broad set of functional and regulatory elements. Enrichment is defined as the proportion of SNP-heritability accounted for by each component divided by the proportion of total SNPs within the module. Components with FDR-corrected enrichment p-values of less than 0.05 were considered significant heritability contributors.

#### Mendelian Randomization

Mendelian Randomization is a form of instrumental variable regression used to formulate a mediating path between a variant (SNP) to a trans-network (trans-eGenes or component individual scores) through a causal gene (cis-gene). We exploit a possible instrument or variant/SNP that changes this causal gene but not the *trans-* gene(s) or component individual scores (aside from through the causal gene). Hence, this yields a mediating path through the casual gene where all the noise is removed except that from the variant. We implement the McDowell, et al. ^34^ formulation of Mendelian randomization.

#### Experimental Validation

Human monocytic cell line, THP-1, was differentiated into macrophages upon treatment with 20 nM phorbol 12-myristate 13-acetate (PMA) for 3 days. Cells were then transfected with human scr-siRNA or CTSB-siRNA (Dharmacon) using Jetprime transfection reagent during 8 h. Cells were collected after 48 h and gene expression levels were assessed by qPCR using Taqman primers.

### Data access

We have made a browser available for all significant *trans*-eQTLs at https://rajlab.shinyapps.io/Tensor_myeloid/. This browser also provides the list of all sparse components, activity scores, and gene scores.

### URLs

LD Score Regression, https://github.com/bulik/ldsc

Sparse Decomposition of Arrays (SDA), https://jmarchini.org/sda/

GWAS Catalog, http://www.ebi.ac.uk/gwas

Myeloid Tensor Shiny Application, https://rajlab.shinyapps.io/Tensor_myeloid/

## Supporting information

## Acknowledgments

We thank B. Fairfax (Oxford University) for sharing the gene expression data. We thank J. Castellano, K. Alasoo, E. Marcora, and members of the Raj Laboratory and Ronald M. Loeb Center for Alzheimer’s disease for critical feedback on the initial draft of the manuscript. This work was supported in part through the computational resources and staff expertise provided by Scientific Computing at the Icahn School of Medicine at Mount Sinai. T.R. is supported by grants from the NIH National Institute on Aging (R01AG054005), the Alzheimer’s Association, and the Michael J Fox Foundation.

## Author Contribution

T.R. conceived the project. S.R. performed statistical and computational analysis with contribution from T.R., E.U., and B.M.S. E.N. and M.P. performed the siRNA experiments. T.R. and S.R. interpreted the data and wrote the manuscript.

