## Supplementary material for "Tensor Decomposition of Stimulated Monocyte and Macrophage Gene Expression Profiles Identifies Neurodegenerative Disease-specific *Trans*-eQTLs"

### Supplementary Figures

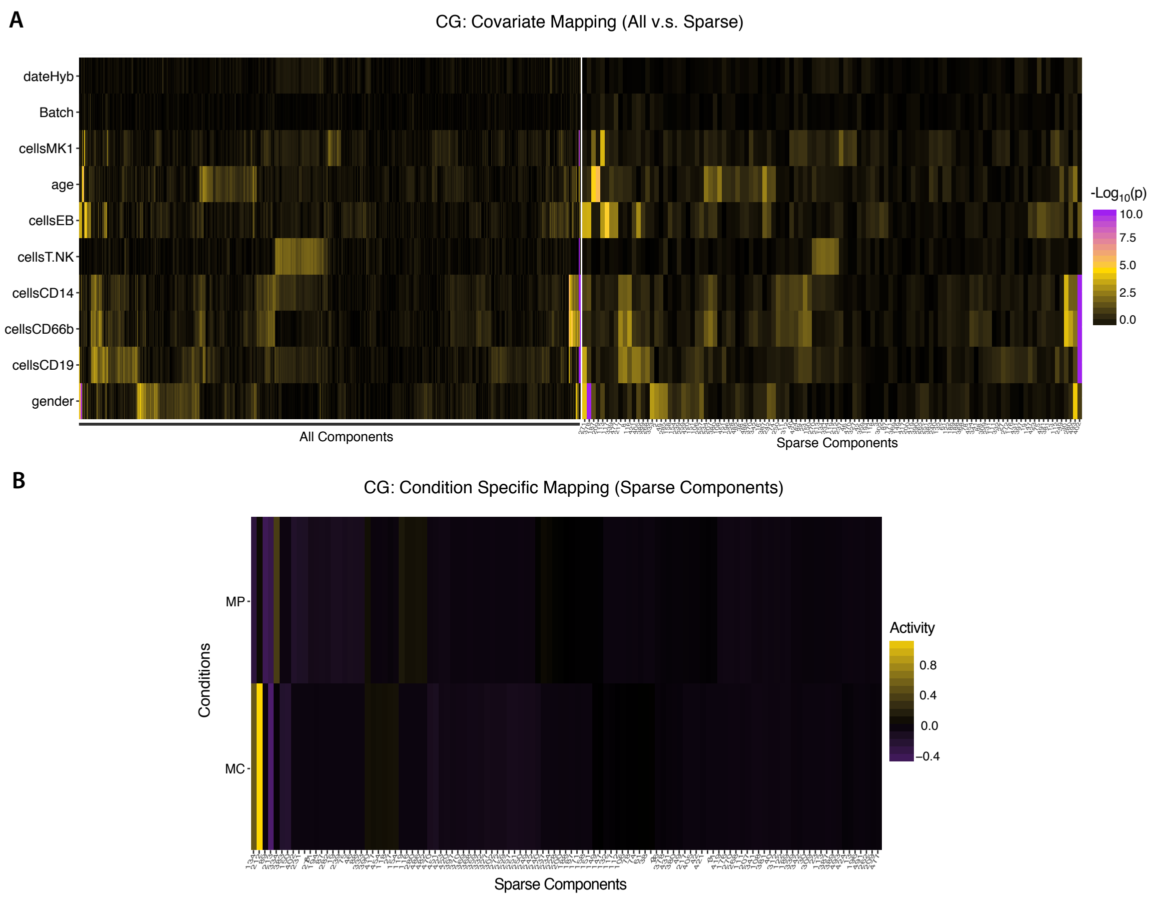


Figure 1: **Association of component individual scores with biological and technical covariates in Cardiogenics (CG).** A: Shown in the heat map are $-log_{10}$ transformed $P$-value of association between CG component individual scores and known covariates. *Left:* All 500 Components, *Right:* Sparse Components. B: For these sparse components, the component tissue scores were grouped based on activity in monocytes (MC) or macrophages (MP).

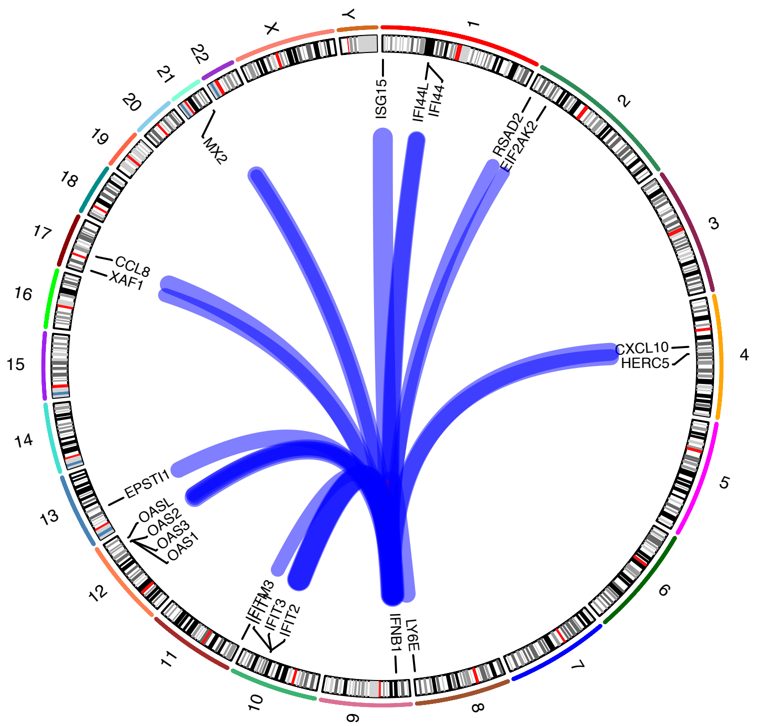


FF: Component 337

Figure 2: **Replication of previously identified *trans*-eQTLs in Fairfax *et. al* (2014).** The SNP rs2275888 maps to FF component 337 enriched with type 1 interferon-related genes with a p-value $= 1.7 \times 10^{-9}$ and FDR $= 1 \times 10^{-3}$ and gene network similar to Fairfax et. al (2014) study. A *cis*-eQTL at rs2275888 for *IFNB1* is associated with the expression of 17 genes in *trans* after 24-hour LPS stimulation, many of which are interferon response genes.

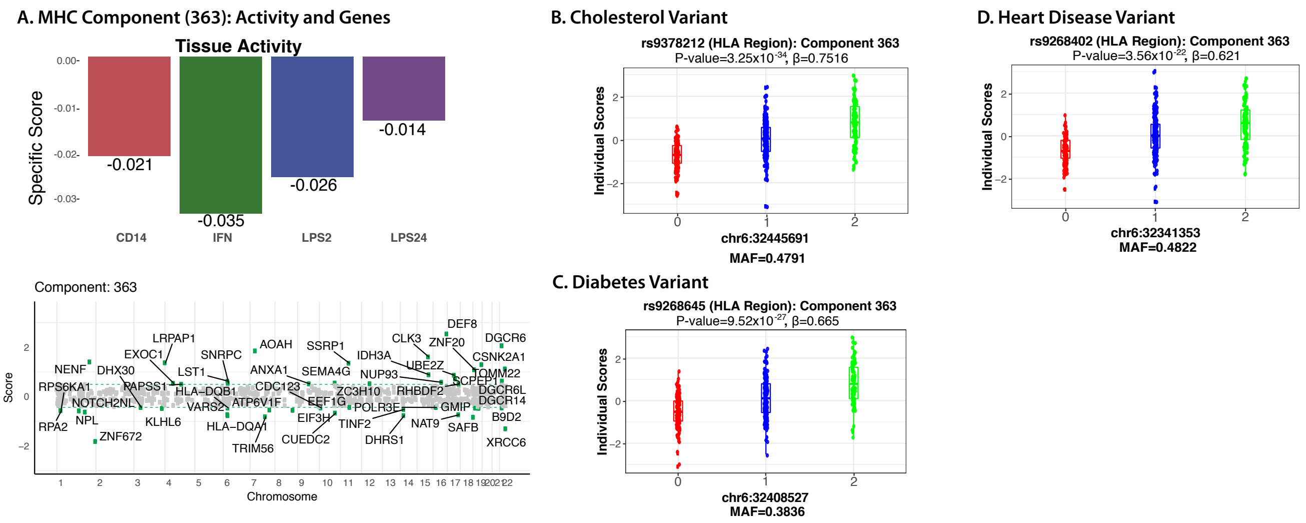


Figure 3: ***Trans*-eQTL mapping to a component with MHC and non-MHC genes.** A: The gene expression profiles of component (363) with MHC and non-MHC genes in FF are shared among all stimuli (Top panel). Show are the gene scores for the genes with Posterior Inclusion Probability (PIP) > 0.5 (bottom panel). *Trans-eQTL* for the MHC component co-localize with B) cholesterol associated risk variant $rs9378212$, C) Type 2 Diabetes risk variant $rs9268645$, and D) Coronary Artery Disease risk variant $rs9268402$.

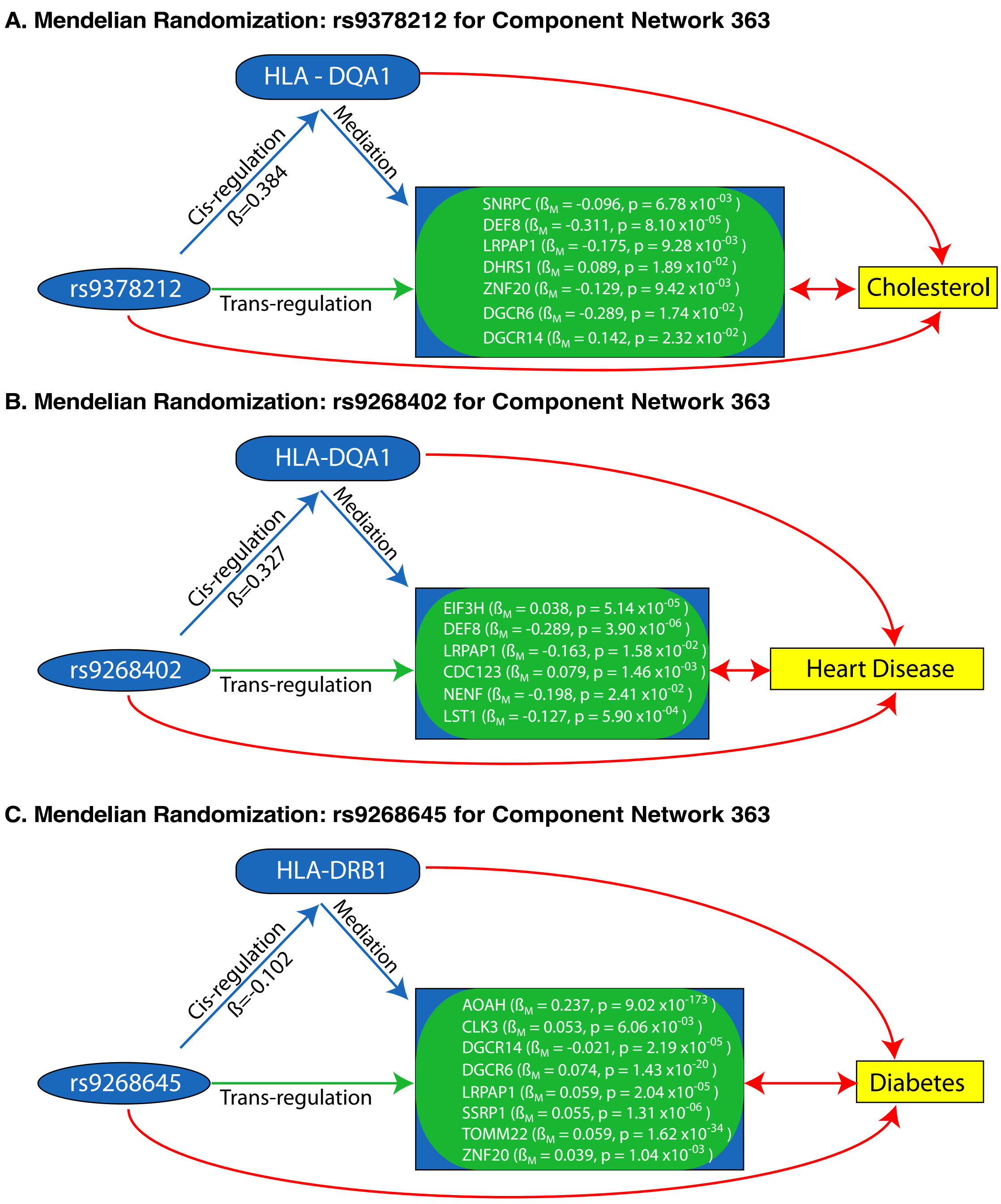


Figure 4: **Mendelian randomization analysis for *Trans*-eQTLs mediated by a *cis*-gene in the MHC.** A: Cholesterol variant *rs*9378212 mediates the *trans*-effects through *HLA-DQA1*, B: Type 2 Diabetes variant *rs*9268645 mediates the *trans*-effects through *HLA-DQA1*, and C: Coronary Artery Disease variant *rs*9268402 mediates the *trans*-effects through HLA-DRB1.

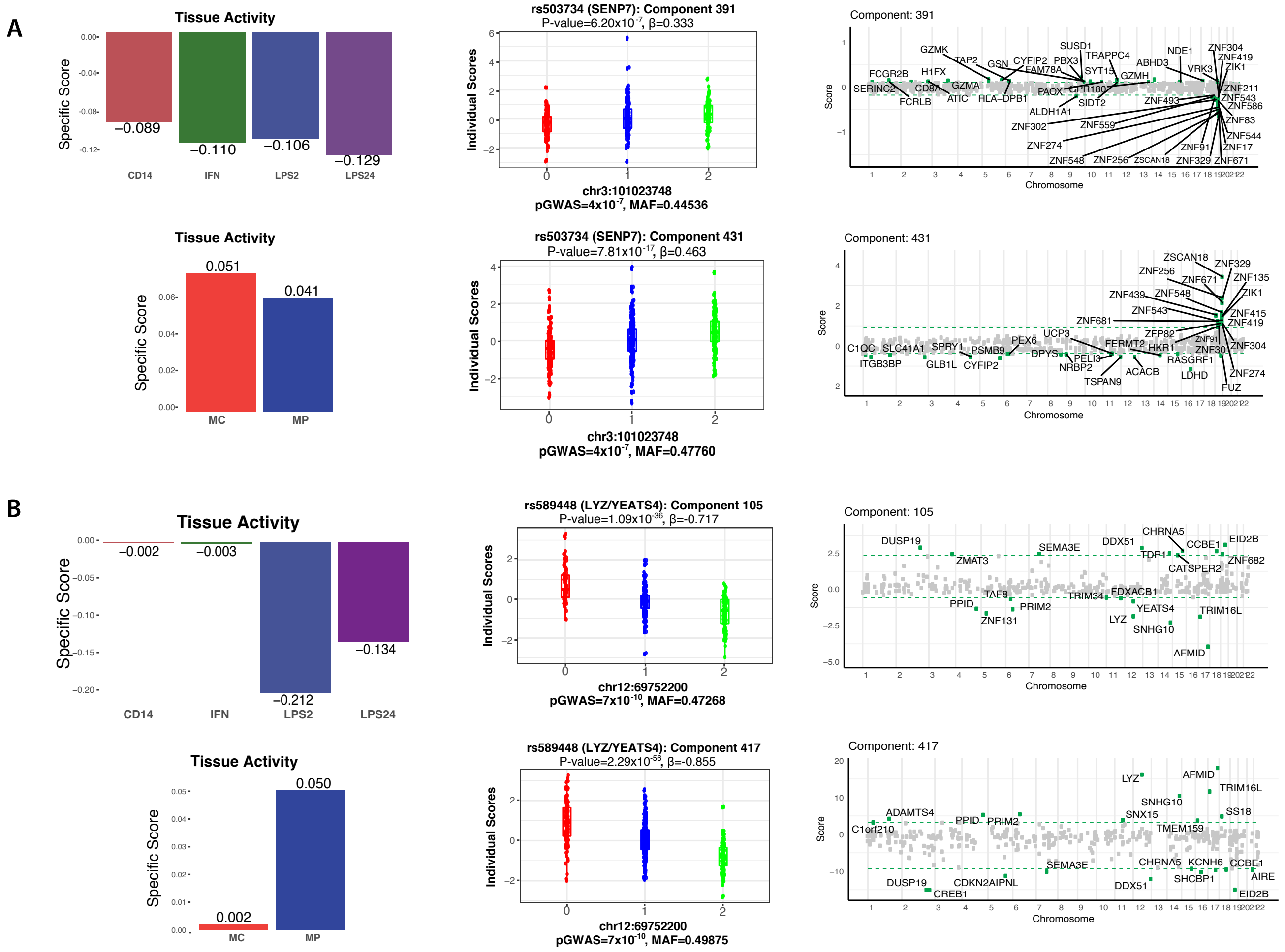


Figure 5: **Replication of *trans*-eQTLs that co-localize with disease-associated susceptibility allele.** A: *Top:* Component 391 active in $FF_{LPS24}$ (left) maps to Crohn's disease associated variant $rs503734$ (Middle). The component contain genes from the members of the Zinc finger family (right). *Bottom:* Replication of the same *trans*-eQTL in $CG$. Component 431 in $CG_{MP}$ (left) maps to the same Crohn's variant (middle). Zinc finger genes shows up in both datasets (right). B: (*Top:*) $FF$ Component 105 active in $FF_{LPS2}$ (left) maps to Coronary Artery Disease variant $rs589448$ (*middle* and ). *Bottom:* The *trans*-eQTL is replicated in $CG$ (Component 417 in $CG_{MP}$ for the same variant $rs589448$ is a *cis*-eQTL to both *LYZ* and *YEATS4* in both FF and CG.

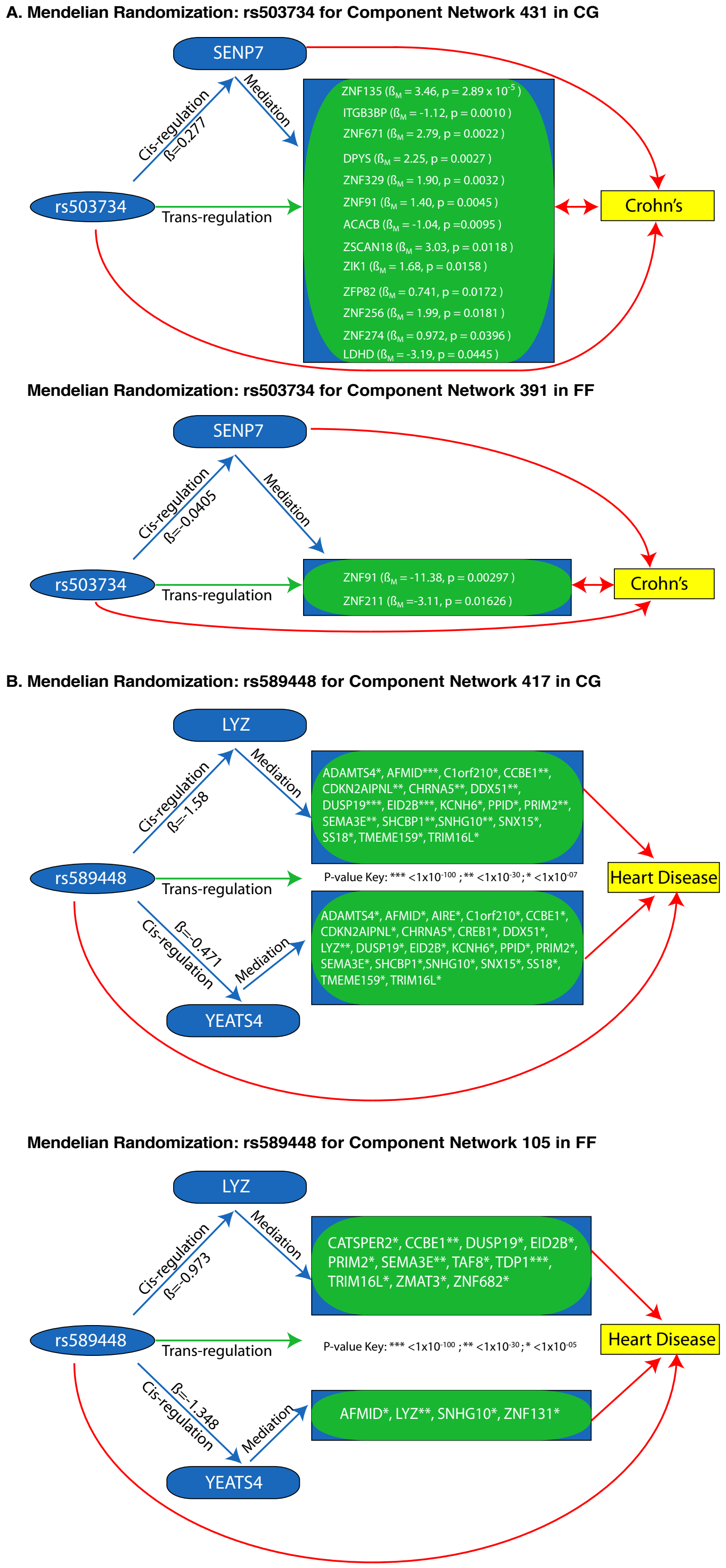


Figure 6: **Mendelian randomization analysis for disease-associated *trans*-eQTLs in FF and CG.** (A) Crohn's variant *rs*503734 mediate *trans*-effects through cis-gene *SENP7* and (B) Coronary Artery Disease variant *rs*589448 mediate *trans*-effects through *cis*-gene *LYZ* and *YEATS4*.

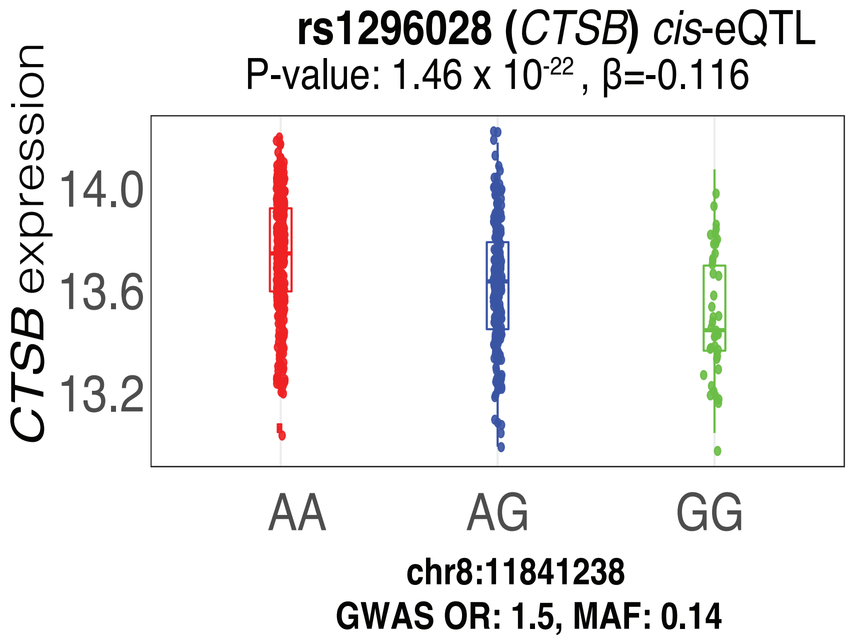


Figure 7: ***Cis*-eQTL** ($rs1296028$-***CTSB***) **co-localizes with Parkinson's disease associated variant** $rs1296028$**.**

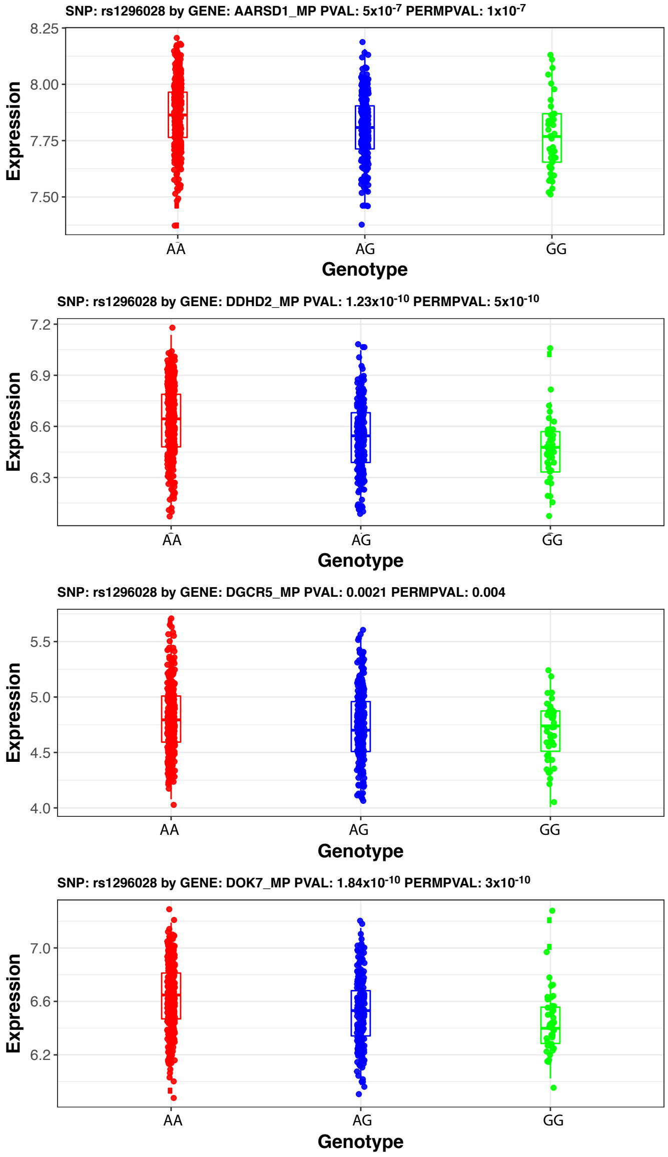


Figure 8: **SNP by Gene *trans*-eQTL association for** *CG* **Component** 46. Shown are box plots for Parkinson's disease associated variant $rs1296028$ (near *CTSB*) mapping to *AARSD1, DDHD2, DGCR5, and DOK7*

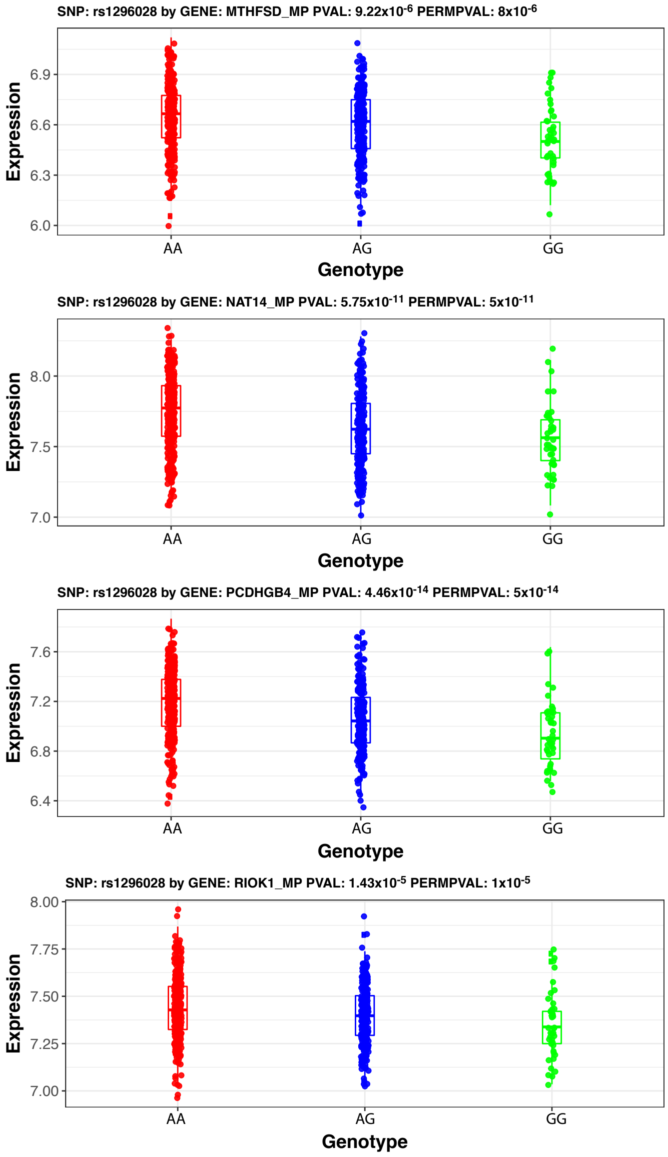


Figure 9: **SNP by Gene *trans*-eQTL association for *CG* Component** 46. Shown are box plots for Parkinson's disease associated variant $rs1296028$ (near *CTSB*) mapping to: *MTHFSD, NAT14, PCDHGB4, and RIOK1.*

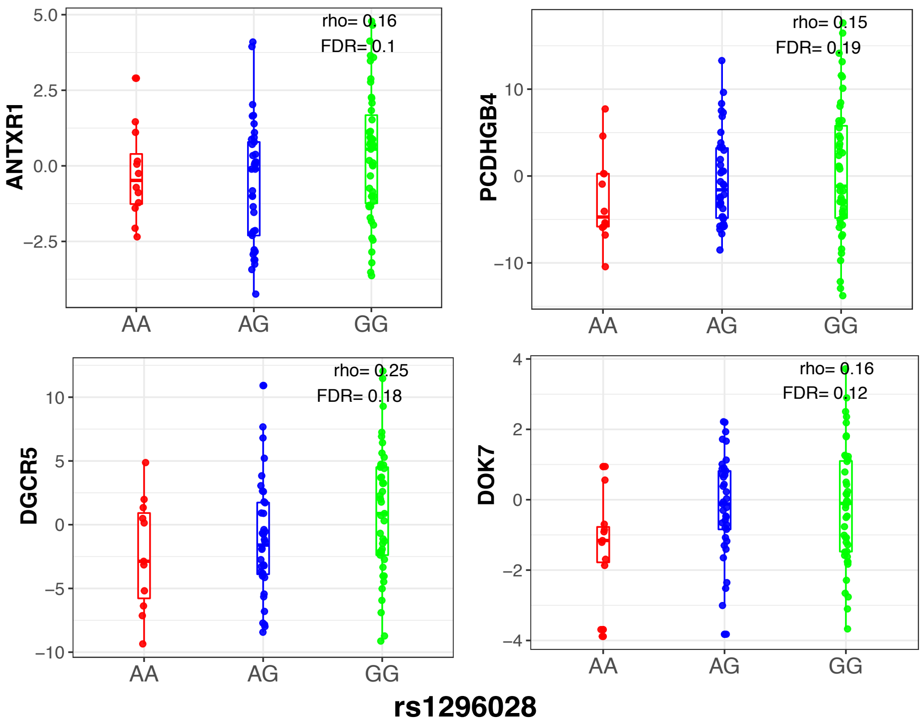


Figure 10: **Independent replication of *trans*-eGenes in the *CTSB* component**. SNP by Gene association analysis was performed in an independent macrophage data from the STARNET cohort. Shown here are the trans-eQTL for rs1296028 and selected *trans* genes in the *CTSB* component at $FDR < 0.20$.

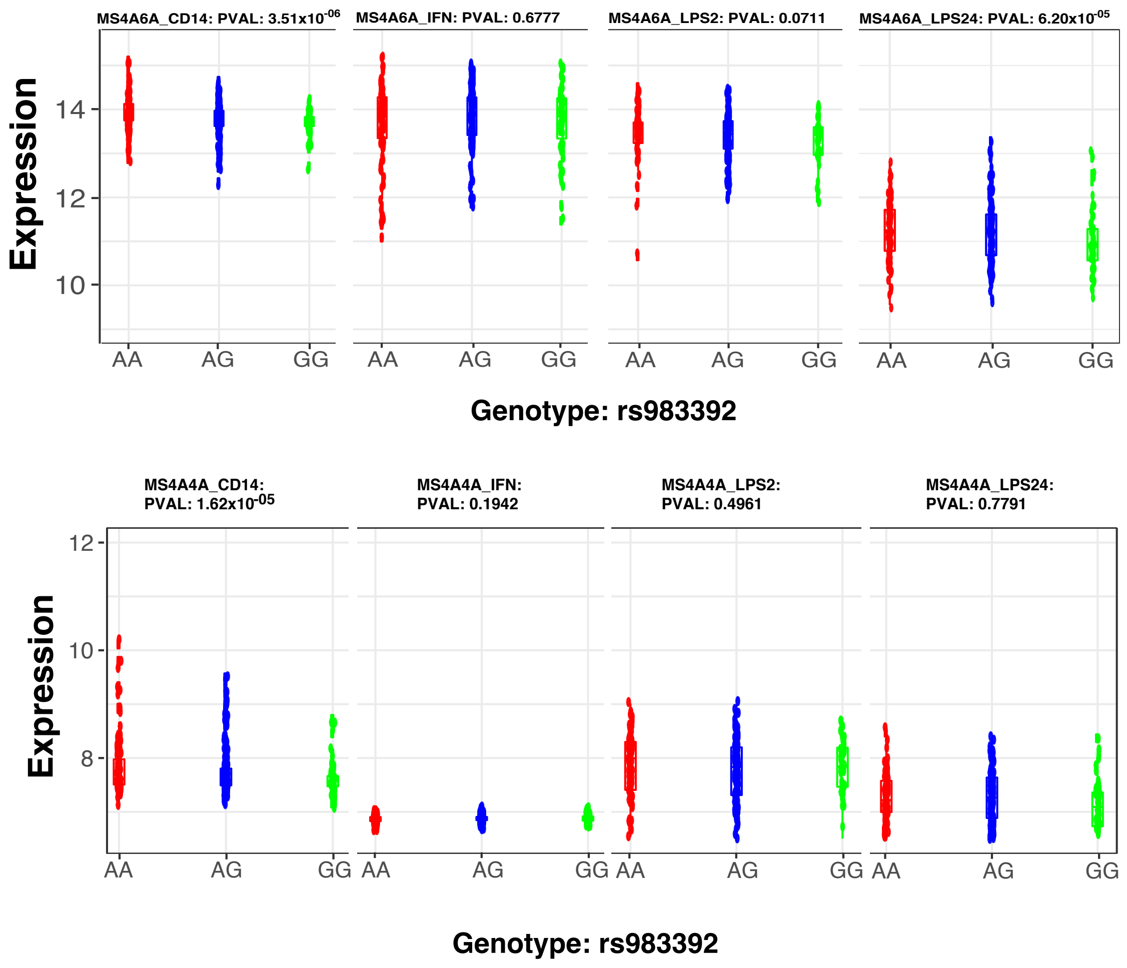


Figure 11: **Significant *cis*-eQTL effect for rs983392 to both *MS4A4A and MS4A6A* in baseline monocytes.**

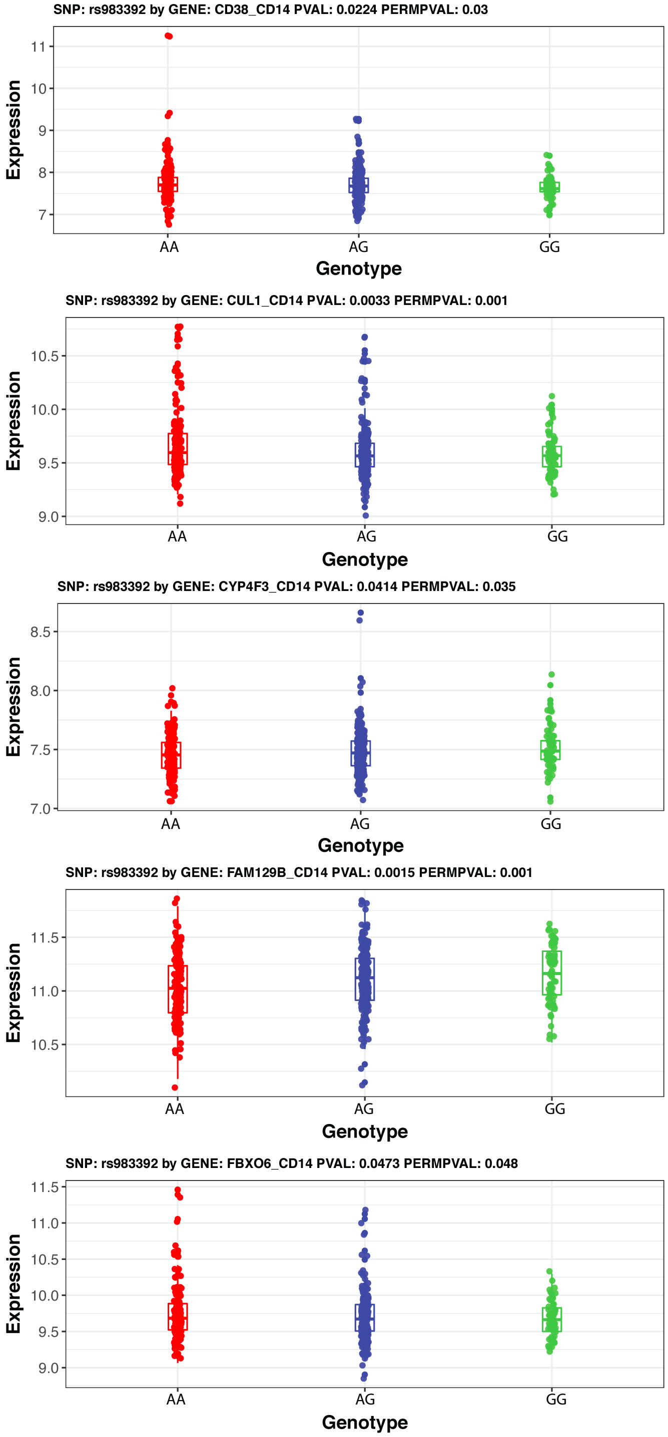


Figure 12: $FF$ **Component 26 trans-eGenes: *CD38, CUL1, CYP4F3, FAM129B, and FBXO6*; SNP by Gene in** $FF_{CD14}$ **for Alzheimer's variant** $rs983392$**.**

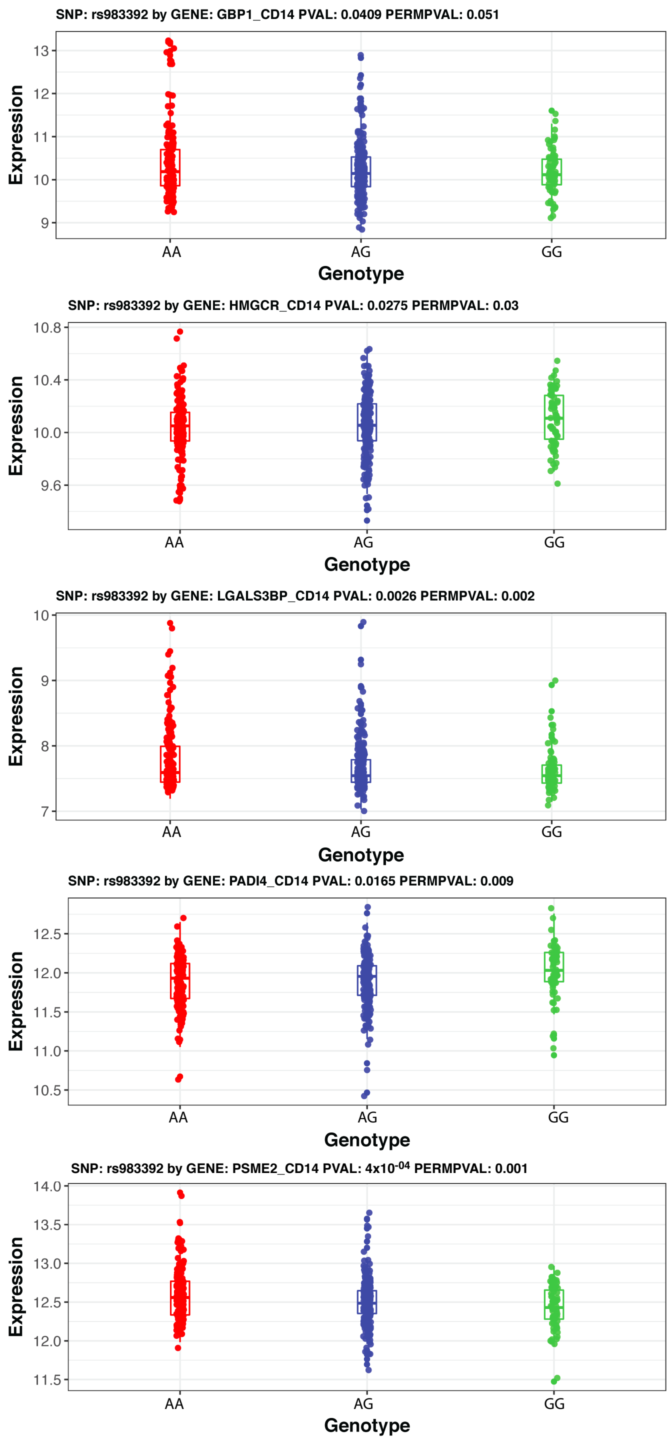


Figure 13: $FF$ **Component 26 trans-eGenes:** ***GBP1, HMGCR, LGALS3BP, PADI4 and PSME2*****; SNP by Gene in** $FF_{CD14}$ **for Alzheimer's variant** $rs983392$

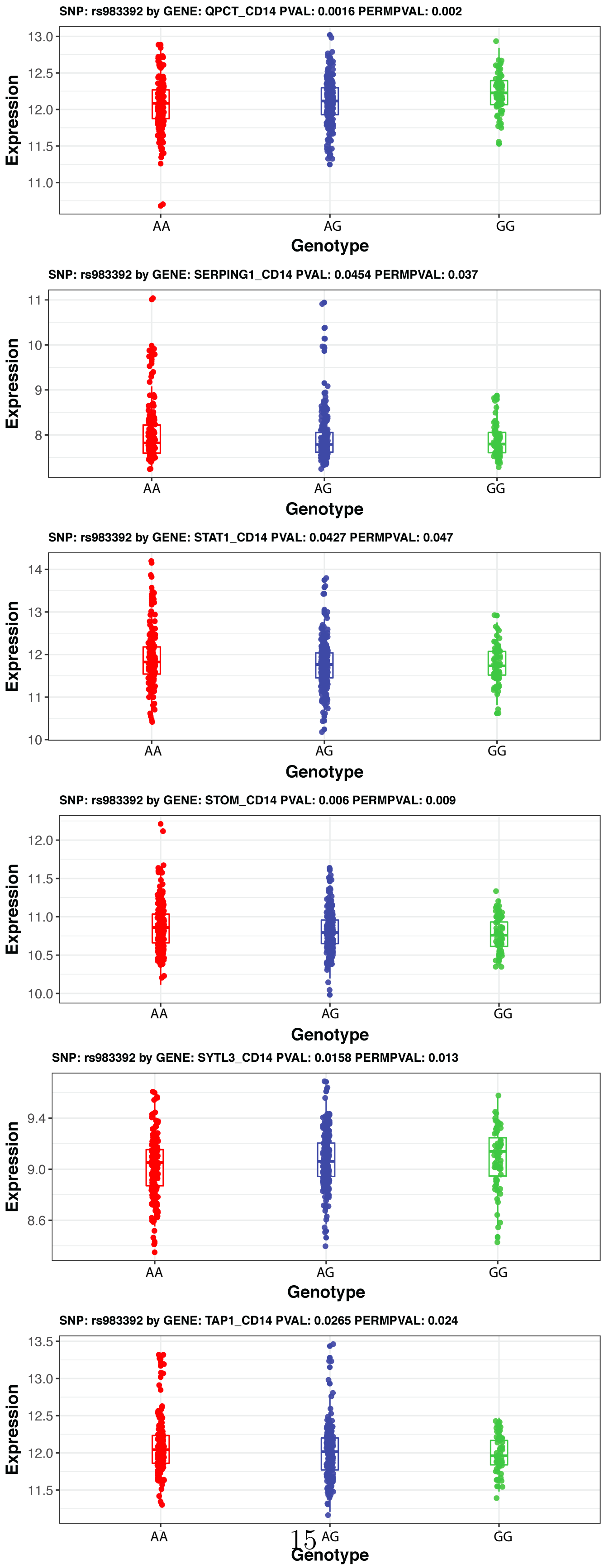




Figure 14: $FF$ **Component** 26 **trans-eGenes:** ***QPCT, SERPING1, STAT1, STOM, SYTL3, and TAP1***; **SNP by Gene in** $FF_{CD14}$ **for Alzheimer's variant** $rs983392$

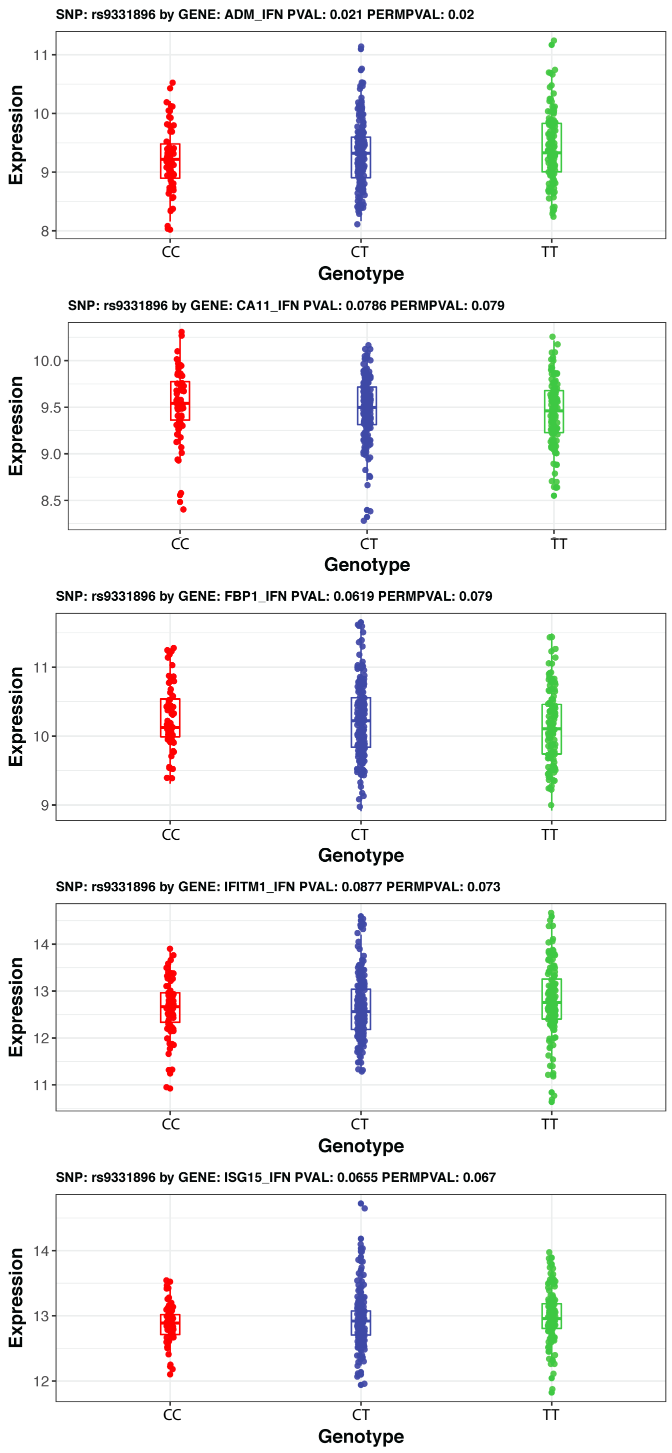


Figure 15: $FF$ **Component** 22 **trans-eGenes: *ADM, CA11, FBP1, IFITM1, and ISG15*; SNP by Gene in** $FF_{IFN}$ **for Alzheimer's variant** $rs9331896$

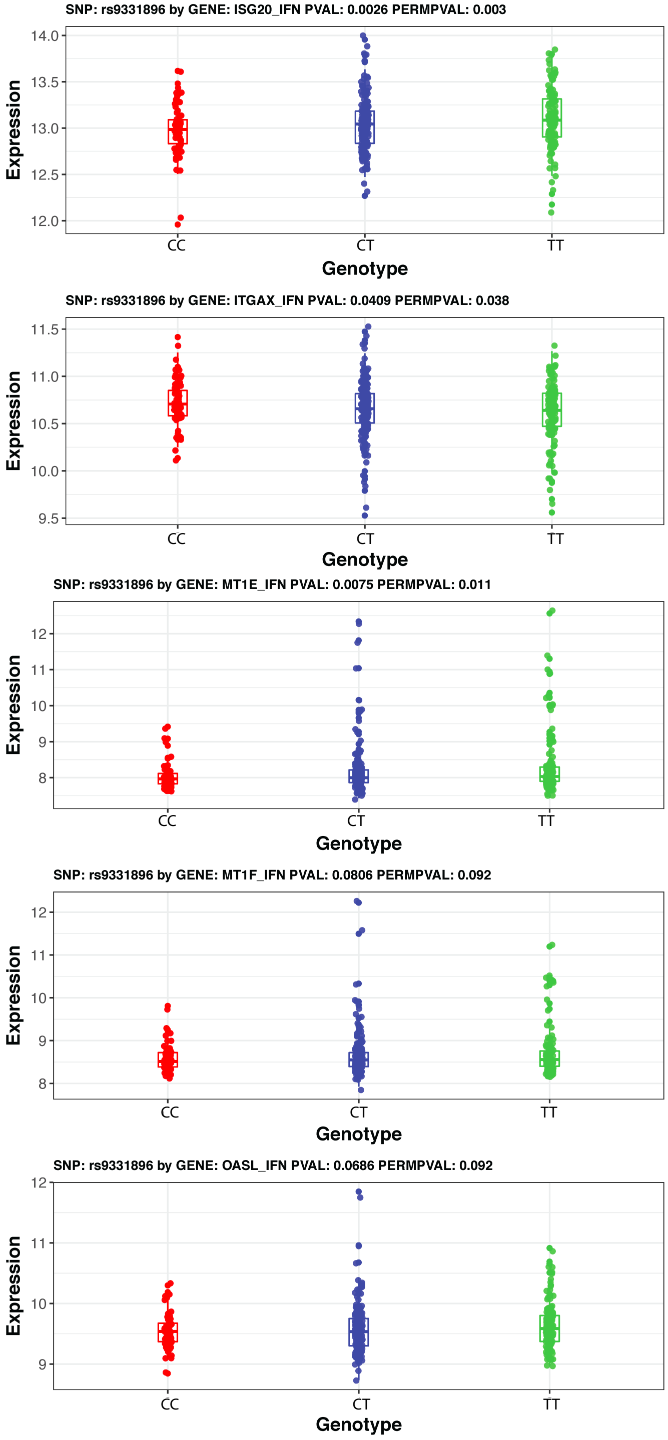


Figure 16: $FF$ **Component** 22 **trans-eGenes:** ***ISG20, ITGAX, MT1E, MT1F, and OASL***; **SNP by Gene in** $FF_{IFN}$ **for Alzheimer's variant** $rs9331896$

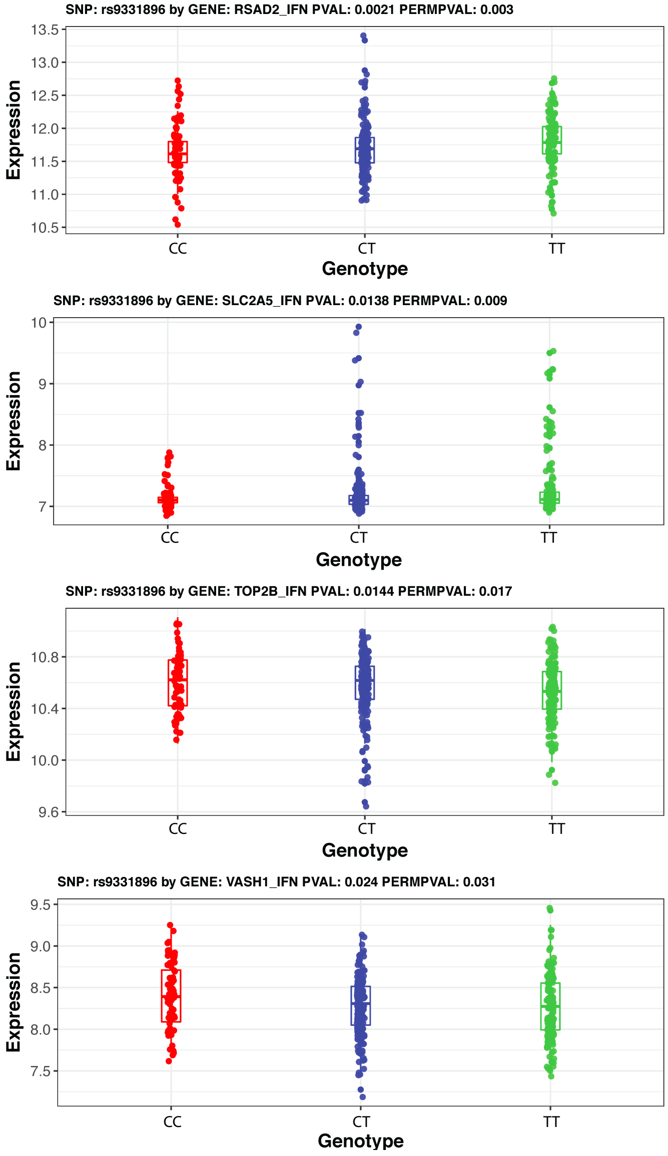


Figure 17: $FF$ **Component 22 trans-eGenes:** ***RSAD2, SLC2A5, TOP2B, and VASH1*****; SNP by Gene in** $FF_{IFN}$ **for Alzheimer's variant** $rs9331896$

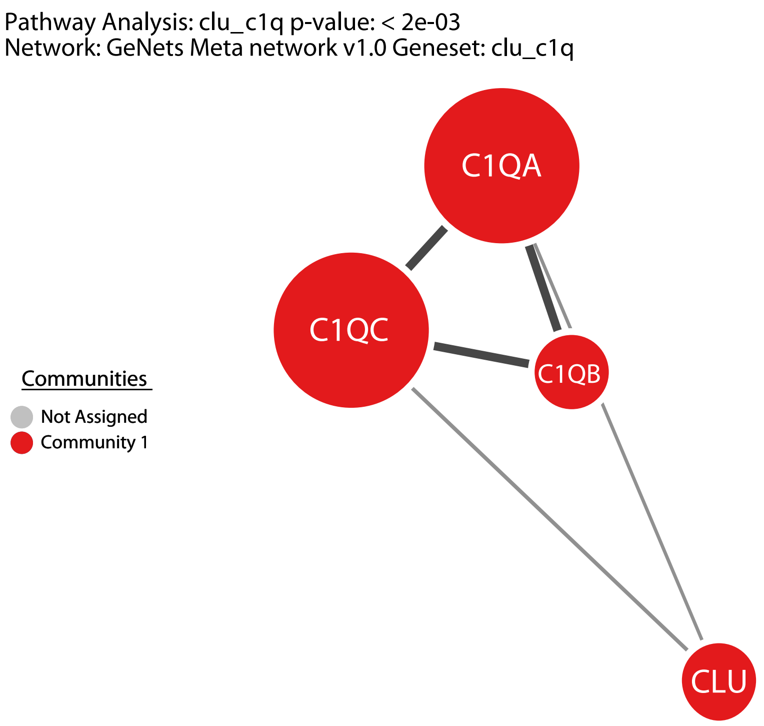


Figure 18: **Protein-protein interaction (PPI) network of protein products of *CLU, C1QA, C1QB and C1QC.*** The PPI suggests that the products of CLU physically interacts with complement proteins.The PPI was generated with GeNets Meta network v1.0 database.

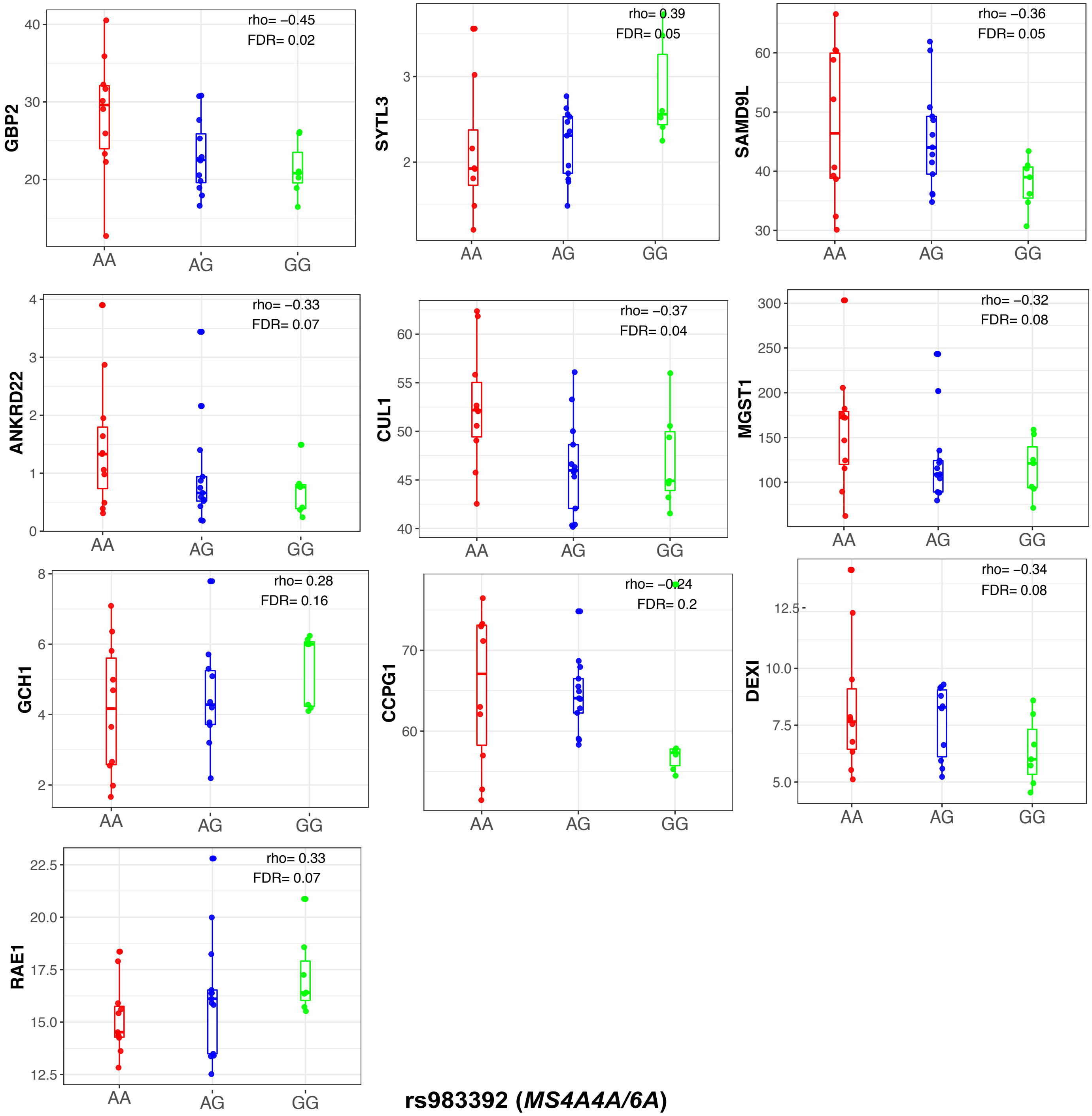


Figure 19: **Independent replication of *trans*-eGenes in the *MS4A4A/6A* component.** SNP by Gene association analysis was performed in an independent baseline monocytes from the ImmVar cohort. Shown here are the *trans*-eQTL for rs983392 and selected *trans* genes in the *MS4A4A/6A* component ($FDR < 0.20$).

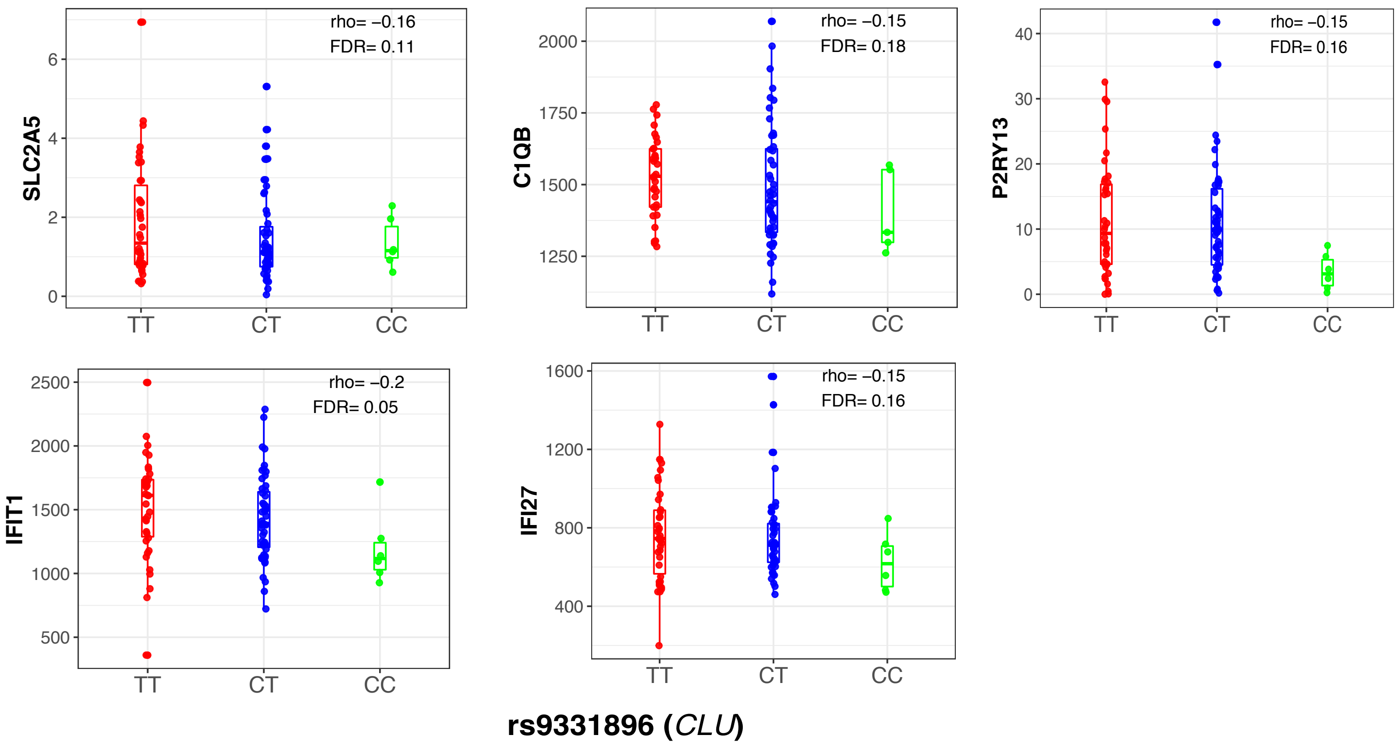


Figure 20: **Independent replication of *trans*-eGenes in the *CLU* component.** SNP by Gene association analysis was performed in an independent stimulated monocytes from the ImmVar cohort. Shown here are the *trans*-eQTL for rs9331896 and selected *trans* genes in the *CLU* component ($FDR < 0.20$).

**Supplementary Tables**

The supplementary tables are available via R Shiny App (see URLs).

Table 1: **List of sparse components with corresponding gene scores and stimuli (IFN and LPS) activity scores identified in the FF dataset.**

Table 2: **List of sparse components with corresponding gene scores and tissue (monocytes and macrophages) activity scores identified in the CG dataset.**

Table 3: **Components that replicate across the three datasets: Fairfax($FF$), Cardiogenics($CG$) and ImmVar($IMM$).** Two-way reverse correlation with the gene scores was conducted on the common intersected genes from the respective components.

Table 4: **Enrichment of Gene Ontology (GO) categories among the genes in the sparse components.**

Table 5: **Sparse components that are enriched for genes within disease-associated loci**. The component number, $P$-value, and corresponding GWAS disease or traits are listed.

Table 6: **SNP-based heritability enrichment for each component.** Proportion of heritability and enrichment statistics for 18 selected complex traits that can be attributed to each sparse component from the FF data.

Table 7: ***Trans*-eQTLs detected in Fairfax data (FF; FDR $< 0.05$).** The matrix-eQTL output with SNP, component number, beta, t-stat, $P$-value and FDR are listed here. Note: The trans-eSNPs are not LD-pruned.

Table 8: ***Trans*-eQTLs detected in Cardiogenics data (CG; FDR $<$ 0.05).** The matrix-eQTL output with SNP, component number, beta, t-stat, $P$-value and FDR are listed here. Note: The trans-eSNPs are not LD-pruned.

Table 9: ***Trans*-eSNPs detected in the FF dataset (FDR $<$ 0.15) that co-localize with trait-associated GWAS SNPs.** The table lists SNP, $P$-value, trait, component number, *cis*-gene (if any), tissue or stimuli specificity scores, genes and gene loading scores for each component and FDR.

Table 10: ***Trans*-eSNPs detected in the CG dataset (FDR $<$ 0.15) that co-localize with trait-associated GWAS SNPs.** The table lists SNP, $P$-value, trait, component number, *cis*-gene (if any), tissue or stimuli specificity scores, genes and gene loading scores for each component, and FDR.

**Supplemental Notes 1**

**1. Assumptions and Sparsity Ranking Statistic Formulation.**

**Implementation assumptions:** In a slight contrast to Hore *et al.,* we impose several strict assumptions prior to implementation. Assumption 1: The unobservable latent space captured by this model is independent for each run. We believe that each model run attempting to capture the latent or underlying space driving the multi-way expression data is independent of each other. Assumption 2: since sparsity is attempted by a spike slab prior driving some values close to (but not necessarily) zero, we use distribution cutoffs to determine thresholding. Since the genes are driven close to but not exactly to zero, top scoring distributional genes are major contributing factor to the network. We use 2.5% tail distributions. Assumption 3: sparsity as per assumption 2 has a maximum limit of T. The idea is that even with distributional thresholding per each component, noisy components may still have a large number of genes. This criterion will reduce their effect of admittance into further analysis. We define T to be 40.

**Ranking Statistic Formulation:** While sparsity is built into the model through the spike-slab prior, low scoring genes in a component are not truly zero but driven close to it. This makes it difficult to determine which component are truly sparse. In an attempt to measure sparsity, a ranking statistic was created. So, for $i = 1, ..., n$, $n$ are number of components, let $R_i(w_i, N_i) = R_i = 1_{\{N_i \leq T\}} \times \dfrac{f(w_i)}{g(N_i)}$ where $w_i$ is a weight function for the significant genes per component based on assumption 3, the functions $f$ and $g$ are mapping functions chosen to reduce the effect of high scoring component gene scores, $N_i$ is the (non-zero) number of significant genes per component and the indicator function drives the ranking of components violating assumption 4 straight to zero. We have defined: $w_i = \dfrac{max_i|x_i| \exp(-(max_i|x_i|)^{-1})}{MAD_i(x_i)}$ where $x_i$ are the actual significant gene scores. This weight function says weight by the largest score proportional to the median absolute deviation (MAD) (assumed to be non-zero) while simultaneously driving small maximum (magnitude) scores close to zero: $f(w_i) = w_i$ and

$g(N_i) = log(N_i)$. The next issue becomes how to choose a cut-off for this ranking statistic (apart from those driven directly to zero). To do this we propose the following cutoff with a bounding proof.

*Proposed cut-off formula and value:* Without attaching distributional assumptions to these functions, we instead propose a formula for cutoff per component then summarize it (through a limiting case). For a (non-zero) median absolute deviation, we let $c$ be the (lower) bound cut-off, then for a given $y = max_i|x_i|$ or maximum absolute score value for that component with a threshold of $N$ ($N_i$ from before) or T (from assumption 3), $c(y, N) \geq \frac{1}{1.4826\exp(y^{-1})log(N)}$. So for the limiting case where $N = T = 40$, $\lim_{y \to \infty} = 0.1828443$ and so our cut-off value is 0.18.
